# Upregulation of Genetic Markers of Poor Prognosis following Chemotherapy in Acute Myeloid Leukemia Cells

**DOI:** 10.64898/2026.02.12.705420

**Authors:** Andrew Taylor, Michael Strasser, Marlee Ng, Irit Rubin, Arja Kaipainen, Angela Oliveira Pisco, Sui Huang

**Author notes:** Divisions of Human Biology and Clinical Research, Fred Hutchinson Cancer Research Center, Seattle, WA, USA. These authors contributed equally to this work (Andrew Taylor, Michael Strasser). **Corresponding Author**: Sui Huang, Institute for Systems Biology, Seattle, WA, USA.

## Abstract

Chemoresistance, a leading cause of treatment failure in cancer, is commonly explained by Darwinian selection of treatment-resistant cell clones. However, recurring resistant tumors invariably display complex phenotypes that contribute to increased malignancy, unlikely to have been selected for by chemotherapy. The growing awareness of (non - genetic) phenotypic plasticity led to the hypothesis that chemotherapy, while reducing tumor burden, also inflicts cell stress that induces stem-like states in cells that survive treatment. Here we examined the transcriptomes of HL-60 leukemic cancer cells that survived exposure to three commonly used drugs at submaximal doses for two to four days, and compared differentially expressed genes to those associated with prognosis in public transcriptome databases of acute myeloid leukemia cohorts. While among the genes differentially upregulated in surviving cells, some reflect the therapeutic effect of chemotherapy as they were associated with favorable outcomes in cohort data, many genes upregulated were associated with poor survival, notably genes involved in stemness, epithelial-mesenchymal transition (EMT), inflammation, drug resistance, and apoptosis evasion. These findings support the idea that treatment effectiveness is the net result of an intrinsic tradeoff: Cytocidal treatment, intended to quantitatively reduce cancer cells, also qualitatively increases the malignancy of non-killed cells, which could contribute to residual disease and relapse. This result has implications for drug screening of new therapeutics, as well as in vitro profiling of patient-derived tumor cell susceptibility to existing drugs, which only assess suppression of cancer cell growth and survival.

**Statement of Significance:** Cells surviving chemotherapy upregulate many genes that may affect prognosis. Therefore, drug screening must embrace a more holistic assessment of the biological quality of cell response beyond the rate of cell killing.

## Introduction

Growing evidence suggests that chemoresistance cannot be explained solely by pre-existing genetic or epigenetic diversity of cancer cells and ensuing selection of variant cells that are constitutively resistant to cytocidal treatment. Instead, following treatment with chemotherapy and radiation, transcriptomic shifts in the non - killed cells triggered by treatment-stress contribute to broad functional phenotype changes (1–5). Non-genetic plasticity of cells, which results from stochastic gene expression, generates substantial phenotypic heterogeneity of cell states within clonal cell populations. When external signals trigger cell state switching, individual cells respond differently, resulting in fractional killing of tumor cells, and altered transcriptomic states in subpopulations of the non-killed cells. In these surviving cells, the cytocidal treatment imparts massive cell stress, and a stress-response consisting of stemness and repair programs is often induced (6–10). This has led to the suggestion that cytocidal treatment is a double-edged sword: While it kills a fraction of cells, hence attenuating tumor burden, it also triggers a repair response in the surviving cell fraction. This stress response is embodied by cell and tissue level programs associated with disease progression: Stemness, inflammation, wound healing, and immune suppression (11–14). Therefore, the dual nature of cancer treatment implies that treatment success hinges on the relative proportions of these opposing responses, and thus, on the nature of stress-induced gene expression programs – a far more complex response than can be captured by the typical readout of “fraction of cells killed at a given dose” used in drug screening.

This recognition led to the idea that genes whose expression are associated with poorer survival can be induced by chemotherapy. This response may underlie the more general phenomenon of treatment-stress induced increase in malignancy: Genes conferring a survival advantage to cancer cells may be upregulated upon treatment, contributing to residual disease and relapse. Thus, we sought to survey this relationship more explicitly and systematically using acute myeloid leukemia (AML) as an example.

We treated HL-60 cells (a well-studied AML cell line) with three different chemotherapy drugs previously studied or currently used in AML treatment (cytarabine, vincristine, venetoclax). We compared the transcriptomes of surviving treated cells to untreated cells to identify genes whose transcript levels increased after chemotherapy treatment. In parallel, we used Cox proportional hazards models to assess the clinical prognosis of gene expression levels measured in untreated patient samples across four different AML cohorts (TCGA AML, Beat AML, AMLCG1, AMLCG2), to identify genes associated with survival. Finally, we assessed the overlap between differentially expressed genes and survival associated genes statistically.

We find a statistical enrichment between “poor prognostic genes” and their downregulation following chemotherapy exposure, reflecting the expected (and desired) consequence of treatment in the form of tumor attenuation, such as differentiation and quiescence (15–18). However, a proclivity towards either malignant or beneficial response to treatment seems to be drug dependent. We also found genes upregulated by treatment that are linked to poor patient survival within our analysis, revealing associations with molecular pathways that promote higher malignancy, such as inflammation, apoptosis evasion, stemness, and epithelial-mesenchymal transition (EMT) related genes. These opposing findings underscore the bifurcation between beneficial response and provoked malignancy that treated cells undergo (19). Transcriptome responses to treatment with cytarabine and vincristine measured at the single-cell level revealed early changes in a substantial portion of HL-60 cells that moved to states similar to those already occupied by just a few rare cells in the untreated cell population, consistent with the theoretical idea (2,20–24) that the apparent induction of a malignant phenotype is not a de novo creation instructed by the treatment, but a cell state shift into latently pre-existing states in gene expression state space facilitated by treatment stress.

## Materials and Methods

### Cell Line, Storage, and Culture

HL-60 cells were [purchased from ATCC](https://www.atcc.org/products/ccl-240) and expanded up to 12 passages, then frozen in Iscove’s Modified Dulbecco’s Medium (IMDM, Invitrogen) supplemented with 20% fetal bovine serum (FBS, Sigma) and 10% DMSO, kept in the vapor phase of liquid nitrogen. Cells from experiments were from passages 16 to 30. These cells were cultured in IMDM with 20% FBS, 1% L-glutamine, penicillin (100 U/mL, Invitrogen) and streptomycin (100 mg/mL, Invitrogen). Cells were maintained at a density between 300,000 and 1,000,000 cells per mL of media and reseeded when the upper density boundary was reached, every 2-3 days. Mycoplasma testing was done upon request by facility staff (last tested September 26^th^, 2024) using [Lonza's Mycoalert Mycoplasma detection assay](https://www.atcc.org/products/ccl-240).

### Live Cell Counts

Live cells were counted using trypan blue stain (.4%, Invitrogen) mixed 1:1 with cell culture suspension and pipetted into cell chamber slides glass slide for live cell counting using Countess FL automated cell counter (Invitrogen).

### Chemicals

The drugs used for in vitro HL-60 treatment and their identifiers are as follows:

vincristine sulfate (referred to as simply “vincristine”, IUPAC: (3aR,3a1R,4R,5S,5aR,10bR)-Methyl 4-acetoxy-3a-ethyl-9-((5S,7S,9S)-5-ethyl-5-hydroxy-9-(methoxycarbonyl)-2,4,5,6,7,8,9,10-octahydro-1H-3,7-methano[1]azacycloundecino[5,4-b]indol-9-yl)-6-formyl-5-hydroxy-8-methoxy-3a,3a1,4,5,5a,6,11,12-octahydro-1H-indolizino[8,1-cd]carbazole-5-carboxylate))
cytarabine (IUPAC: 4-amino-1-[(2R,3S,4S,5R)-3,4-dihydroxy-5-(hydroxymethyl)oxolan-2-yl] pyrimidin-2-one)
venetoclax (IUPAC: 4-(4-{[2-(4-Chlorophenyl)-4,4-dimethyl-1-cyclohexen-1-yl]methyl}-1-piperazinyl)-N-({3-nitro-4-[(tetrahydro-2H-pyran-4-ylmethyl)amino]phenyl}sulfonyl)-2-(1H-pyrrolo[2,3-b]pyridin-5-yloxy)benzamide)

### Drug Treatment, FACS and Sequencing

HL-60 were administered chemotherapy at half maximal effective concentration (EC50) doses or higher over 48-96 hours, depending on experiment, to impart massive stress to cell populations while still leaving a portion of cells alive for sorting and RNA sequencing.

Bulk RNA-seq was performed using the Illumina NovaSeq 6000 platform with PE150 reads. Cells were plated at a density of 200,000 cells/mL and treated the next day with the following drug concentrations, all above EC50: cytarabine 5 μM, venetoclax 500 nM, vincristine 5 nM. Cells were sorted using a BD Influx flow cytometer (BD Biosciences) on day three (vincristine) or four (cytarabine and venetoclax). Mock treated cells (i.e. cells treated with an equal volume of drug carrier solution) were sorted alongside drug-treated cells. Cells were gated for no/low propidium iodide (PI) and immediately resuspended in TRIzol LS for sample preservation prior to RNA extraction and sequencing library preparation for sequencing.

Single-cell RNA-seq for “Experiment A” was done using platform Chromium Next GEM Single Cell 3’ Reagent Kit v3.1, followed by library construction and Illumina sequencing. HL-60 cells were plated at a density of 400,000 cells/ml. Cells treated with cytarabine were given 1.6 μM (precisely EC50 concentration), and cells treated with vincristine were given 5nM. Live cells were harvested by sorting for no/low PI using fluorescence-activated cell sorting (FACS) and then fixed in 80% methanol for sequencing on d0 and d2 of treatment.

Single-cell RNA-seq for “Experiment B” was performed using the Chromium Single Cell 3’ Reagent Kit V2, followed by sequencing library construction and Illumina sequencing. HL-60 cells were plated at a density of 400,000 cells per mL with 5 nM vincristine, with live cells harvested by sorting for no/low PI using FACS on each day of treatment (d0, d1, d2) and frozen in 10% dimethyl sulfoxide (DMSO) prior to sequencing.

### Patient Cohort Data Processing and Exclusion Criteria

Beat (version 1.0) (25), TCGA (26) and German AML Cooperative Group (AMLCG) 1999 leukemia trial (27) data were retrieved as specified in the “data availability” section.

Deposited gene expression microarray data for AMLCG was already Robust Multichip Average (RMA) (28) normalized and QC filtered as described in (29). The treatment regimen for AMLCG patients is described here (30), with limited clinical features (age and M3 classification) available from GEO accession. Patients in AMLCG Cohort 1 (n=422) were those analyzed in parallel using two partially overlapping (but largely distinct) probe sets, HG-U133A and HG-U133B. For probes measured twice, only values from probe set A were considered and set B was excluded. Patients in AMLCG Cohort 2 (n=140) were those analyzed using microarray probe set HG-U133 Plus 2.0. In cases where microarray probe to gene mappings in the AMLCG cohorts 1 and 2 were not unique, probes mapping to several genes were excluded and genes mapping to several probes had their probe expression levels averaged.

Transcript per million (TPM) gene expression data obtained from Beat and TCGA were log2(x+1) transformed (microarray data was not, as log2 scaling is already done during RMA normalization).

Gene expression values for each cohort were then z-scored across patient samples within the cohort prior to Cox survival regression, such that results would become comparable between genes and across cohorts. That is, for each gene in a given cohort, its hazard ratio was defined in terms of standard deviations from the cohort mean. Genes with extremely low variance (less than .0001) were omitted from testing and marked as “not prognosis associated”.

Patients under the age of 18 or documented as M3/ Acute Promyelocytic Leukemia (APL) / PML-RARA fusion were omitted on the basis of phenotypic differences with most adult AMLs that can lead to erroneous interpretations during survival analyses. Of further note, patients receiving non-standard treatment (targeted or palliative) or bone marrow transplants were removed from the Beat and TCGA cohorts in the interest of interrogating response to standard chemotherapy treatments while also keeping initial transcriptomic measurement tightly coupled to survival data. AMLCG cohorts were not subject to bone marrow transplant restrictions due to missing information. Samples collected using leukapheresis (only blood or bone marrow aspirate allowed), or from relapsed patients were also excluded from analysis (seen in Beat and TCGA data).

### Patient Probe and Gene ID Conversion

Following survival regression, microarray probe IDs used in the AMLCG cohorts were converted to ENSG identifiers (Beat and TCGA gene IDs were already provided as such). ENSG identifiers from all cohorts were then further converted to HGNC/ HUGO nomenclature for visualization and result reporting. This was done using BioMart (https://www.ensembl.org/biomart).

### Survival Regression

To determine poor prognostic markers, genes measured in each cohort (at diagnosis) were analyzed iteratively under two different covariate inclusion schemes: univariate (gene only), and multivariate (gene, age, white blood cell count in trillions/liter) to adjust for potential mediators or confounders.

AMLCG cohort data did not include white blood cell count, and thus white blood cell was not be used as a covariate for multivariate regression (just age and gene expression value).

In the case that additional covariates were missing for multivariate analysis (other than white blood cell count for AMLCG patients), patients were omitted from this analysis.

Analysis was done using Cox proportional hazards regression in the Python package “Lifelines” v0.29.0 (31). The baseline hazard was estimated using Breslow’s method. Parameters were fit using the Newton-Raphson algorithm and p values were estimated using the Wald test (as specified in “Lifelines” code base and documentation). Multiple testing correction (Benjamini-Hochberg, BH) was applied to all genes tested.

### Differential Expression Analysis (Bulk Sequenced HL-60 Drug Treatments)

Bulk RNA-seq reads were pseudoaligned to the GRCh38 reference transcriptome (ENSEMBL release 96) with kallisto (v.0.48) using the default k-mer size=31 (32). Differential expression between controls and treatment was estimated using the package DESeq2 (with experiment batch as an additional covariate) and fold changes were shrunk via “apeglm” (33). Log2 fold change (log2FC) was reported, and false discovery rate (FDR) was calculated using BH multiple testing correction (34).

### Single-Cell Analysis (Leiden Clustering, Cluster Marker Genes, Cell Cycle Inference)

Single-cell RNA-seq reads were pseudoaligned to the GRCh38 reference transcriptome (ENSEMBL release 96) with kallisto (v.0.48) using the default kmer size=31 and transformed into a cell-by-gene count matrix with bustools (v.0.40) (35). Cell barcodes were filtered according to the 10x Genomics whitelist (v3).

Using Scanpy (v 1.10.2) (36), single-cell RNA-seq data was preprocessed, cell count normalized, log transformed, genes z-scored, and filtered for highly variable genes for principal component (PC) analysis. The first 50 PCs were used to construct a K-nearest neighbors graph and perform Leiden clustering. Cell cycle inference based on cell cycle phase specific gene marker expression was done using Scanpy’s internal function. The above were done separately for both single-cell RNA-seq vincristine experiments. In each instance, marker genes were determined using one vs. all other cluster Welch’s t-tests. Cluster similarity was computed across experiments using marker genes for each cluster (top 100 sorted by FDR, provided FDR < 0.05 and log2FC > 0.7) by calculating the Jaccard index.

### Gene Ontology Enrichment

Gene ontology enrichment was performed using the GSEApy (v 1.1) Python wrapper for Enrichr (37), which assessed enrichment between associations in curated gene sets “GO_Biological_Process_2023”, “KEGG_2021_Human”, and “MSigDB_Hallmark_2020” and gene lists assessed using Fisher’s exact test (for overrepresentation/ enrichment in 2x2 contingency tables). Gene ontology enrichment was done for the intersections of upregulated or downregulated genes across drug treatments (bulk sequenced) with curated genes sets. Similarly, this was also done for the intersections of poor or favorable prognosis associated genes across leukemia cohorts with curated gene sets. Multiple testing correction (BH) was applied within each curated gene set, as each set contains many biological processes/ associations tested for enrichment).

### Enrichment Testing between Prognostic and Differentially Expressed Genes

Fisher’s exact test for 3x3 contingency tables was used to initially check for an association between genes with associations of poor, favorable, or no discernible prognosis and genes that were upregulated, downregulated or not discernibly differentially expressed. This was done using the base “stats” package in R with its implementation of Fisher’s exact tests for tables greater than 2x2, using network algorithms to arrive at an approximate p value (38,39).

Post-hoc testing for specific associations involved collapsing the aforementioned 3x3 tables to 2x2 tables, such that rows and columns not of interest for a given test were merged. For instance, if our association of interest was between poor prognostic and upregulated genes, then rows corresponding to no prognosis and favorable prognosis were collapsed into a single row, just as columns corresponding to no differential expression and downregulation were collapsed into a single column, effectively creating a table with rows of “upregulated” and “not upregulated”, and columns of “poor prognostic” and “not poor prognostic”. This was done iteratively such that all four combinations of poor/upregulated, poor/downregulated, favorable/upregulated, favorable/downregulated could be assessed for enrichment after for a given 3x3 table. This process of collapsing tables to assess more specific enrichments was then repeated for all pairwise combinations of our three drug perturbations (differential expression results are drug specific) and four cohorts (prognosis associations are cohort specific).

Importantly, 2x2 tables were specifically tested for enrichment using Fisher’s exact test for overrepresentation (a specific “one-sided” enrichment test, not possible in 3x3 cases), as defined by an overabundance of genes occupying contingency table intersections of interest beyond what is expected at random, rather than a depletion, or two-sided tests which would capture either case (i.e. a lack of independence between categories. This corresponds to setting the “alternative” parameter to “greater” in the [SciPy Python implementation of Fisher's Exact Test](https://docs.scipy.org/doc/scipy/reference/generated/scipy.stats.fisher_exact.html), where the exact calculation is further explained.

### ChEA3 Transcription Factor Inference

Genes found to be poor prognostic in any cohort and upregulated following at least two out of three drug treatments (where surviving cells were bulk-sequenced) were also queried using ChIP-X Enrichment Analysis Version 3 (ChEA3) (https://maayanlab.cloud/chea3/) (40) to infer transcription factors (TFs) regulating these genes. TFs were assigned composite scores by ChEA3 (Mean Rank and Top Rank) by integrating results across orthogonal “omics” datasets as described in the release paper. We selected mean rank scores for each TF as our inference metric, whereby lower numerical scores indicate TFs that are more highly inferred. The downloadable results TSV file available upon query contained specific correspondences between queried genes and their inferred transcription factors, which were used to generate an adjacency matrix representing interactions between select genes and their top 20 inferred TFs, clustered among regulated genes to indicate regulons (see results).

### Data Availability

In vitro chemotherapy treatment data generated in this study will be made **available during peer review and upon publication.**

Beat (version 1.0) (25) and TCGA (26) cohort gene expression data (TPM) were retrieved from GDC data portal download tool (https://portal.gdc.cancer.gov under data release v40.

Clinical and survival data were obtained as follows: Beat AML information was accessed from the Vizome website “Clinical Summary” link (https://biodev.github.io/BeatAML2) and updated TCGA AML clinical and survival data were queried from https://gdc.cancer.gov/about-data/publications/laml_2012 under “updated supplemental table 01”.

German AML Cooperative Group (AMLCG) 1999 leukemia trial data, including gene expression, clinical, and survival data, was downloaded from Gene Expression Omnibus database (GEO) accession GSE37642.

Analysis code is available at github.com/andrew-taylor-github/upreg-markers-poor-prog-following-chemo-aml

### Generative Artificial Intelligence

ChatGPT was used to generate occasional code snippets and conduct more tailored literature searches.

## Results

### HL-60 Drug Treatment Experiments: Determining Drug-Response Genes

To determine the molecular changes induced by chemotherapy treatment in the non - killed cancer cells, HL-60 cell cultures were treated with a single drug (vincristine, venetoclax or cytarabine) at submaximal doses just above EC50 concentrations, or mock-treated, with 3 biological replicates per condition, for 72 hours (vincristine) or 96 hours (cytarabine and venetoclax). Surviving cells were sorted by FACS based on propidium iodide staining of dead cells, and live cells collected for bulk RNA sequencing, yielding a total of 12 samples (4 conditions, 3 replicates). Principal component analysis **(Supp. Fig. 1A)** shows clear separation of control and drug-treated samples. We first determined differentially expressed genes (see methods) between control cultures and each of the three drug-treated cell cultures (FDR< 0.05 and |log2FC| > 0.7, **Supp. Fig. 1B**). We found 5723 upregulated genes (with respect to control) in the non-killed cells after treatment with cytarabine, and, 841 and 935 upregulated genes for vincristine and venetoclax, respectively. A common set of 490 genes was upregulated by all three drugs (**Supp. Fig. 1C**). Similarly, we found 4128 downregulated genes after treatment with cytarabine, and 341 and 575 downregulated genes for vincristine and venetoclax, respectively. Here, the common set contained 166 genes downregulated by all three drugs (**Supp. Fig. 1E**). Gene ontology analysis of the genes upregulated in all three drugs (**Supp. Fig. 1D**) using Enrichr indicated hypoxic, xenobiotic and inflammatory stress responses mediated in part by cytokine, toll-like receptor, and NF-kB signaling, as the top results. Similar analysis of genes downregulated across all three drugs suggested changes to biosynthetic processes and metabolism (**Supp. Fig. 1F**).

### Survival Analysis in AML Cohorts: Determining “Prognostic” Genes

Associations between gene expression levels (measured in AML patients prior to receiving treatment) and survival times were determined in publicly available AML transcriptome cohort data, using Cox proportional hazards regression to fit each gene independently, first with no other covariates (univariate). Poor prognostic genes were defined as having a positive log hazard ratio, and conversely, favorable prognostic genes as having a negative log hazard ratio (provided FDR < 0.05). Hazard ratios and FDRs are plotted in **Supp. Fig. 2A**. The AMLCG1, AMLCG2 and Beat cohorts returned 150, 51, and 219 poor prognostic genes respectively, with six genes overlapping between three cohorts (**Supp. Fig. 2B**). The TCGA cohort yielded no statistically significant prognostic genes, likely due to patient exclusion criteria (see methods) that led to a small cohort size (n=58) compared to AMLCG1 (n=398), AMLCG2 (n=129), and Beat (n=141). Inspection of favorable prognostic genes showed only two genes shared between the AMLCG1 and Beat cohorts, with none shared across any cohorts otherwise (**Supp. Fig. 2D**). The Beat cohort had only six genes associated with favorable prognosis, while analysis of AMLCG1 and AMLCG2 yielded 67 and 40 favorable prognosis associated genes respectively.

This analysis was extended to the multivariate form (**Supp. Fig. 3**) where age, and white blood cell count (available in Beat and TCGA but not AMLCG), were included as covariates during regression, yielding fewer prognostic genes but showing similarity to previous univariate results. Patients with missing covariate values were removed from analyses including those covariates.

### Combining Drug Response and Survival Data

To determine if there is a statistical dependence between the two gene properties “induced by treatment” and “associated with survival” for each drug and cohort pair, we constructed 3x3 contingency tables with columns accounting for the directionality of differential expression (genes upregulated by treatment, downregulated, or neither) and rows corresponding to survival association (genes associated with poor survival outcome, favorable, or neither). This resulted in a set of 12 contingency tables (3 drugs x 4 cohorts) for the case where gene-prognosis associations were determined using univariate survival regression (gene only) and another set of 12 tables for multivariate regression (gene and other clinical covariates).

All contingency tables were then assessed using Fisher’s exact test for 3x3 tables (see methods) to (i) determine if there was a general statistical dependence between rows (prognosis) and columns (differential expression due to treatment) and (ii) perform a robust analysis that eliminates idiosyncrasies of a given drug, cohort or covariate choice in survival regression. This analysis reveals that a gene’s differential expression following in-vitro chemotherapy treatment is generally not independent (p value < 0.05, Fisher’s exact test) of its association with survival, observed in all contingency tables other than those constructed using prognostic associations from the TCGA cohort (**Supp. Fig. 4**), where enrichment is impossible as no genes reached the FDR value required to be considered prognostic.

### Enrichment for Category Overlaps

Because Fisher’s exact test in 3x3 tables (or any table larger than 2x2) cannot reveal the exact nature of the dependence (e.g., whether upregulated genes are more frequently associated with poor survival than expected by chance), further post-hoc tests (Fisher’s exact tests for overrepresentation) were performed to study the connection between drug-induced genes and survival in more detail. This was done by collapsing our original 3x3 tables into 2x2 tables (described in **Supp. Fig. 5** and methods) in such a way that we could assess enrichment for four different overlaps of interest: “poor prognostic and upregulated”, “poor prognostic and downregulated”, “favorable prognostic and upregulated” and “favorable prognostic and downregulated”. Importantly, here enrichment (overrepresentation) of specific overlaps is defined as an overabundance of genes occupying the specific contingency table intersections of interest, beyond what is expected at random; this is in contrast to testing for depletion, or two-sided tests which would capture either case.

Among these intersections, an over representation of genes meeting the criteria of both “upregulated” and “poor prognostic”, or “downregulated” and “favorable prognostic”, could be interpreted as supporting the emerging notion that treatment-stress can, in view of cell plasticity, trigger a shift towards higher malignancy (e.g. the upregulation of an oncogene or the downregulation of a tumor suppressor). We term this behavior “malignant response”. By contrast, genes that belong to the categories of significantly “downregulated” and “poor prognostic”, or “upregulated” and “favorable prognostic”, might reflect the response to effective treatment in the non-killed cells, such as cellular differentiation and cell cycle arrest. We term this “beneficial response”. To better understand associations between drug responses and these outcomes, we once more iterated through our cohort and drug selections, performing these more granular 2x2 Fisher’s exact tests for overrepresentation (**Fig. 1**).

**Figure 1:**
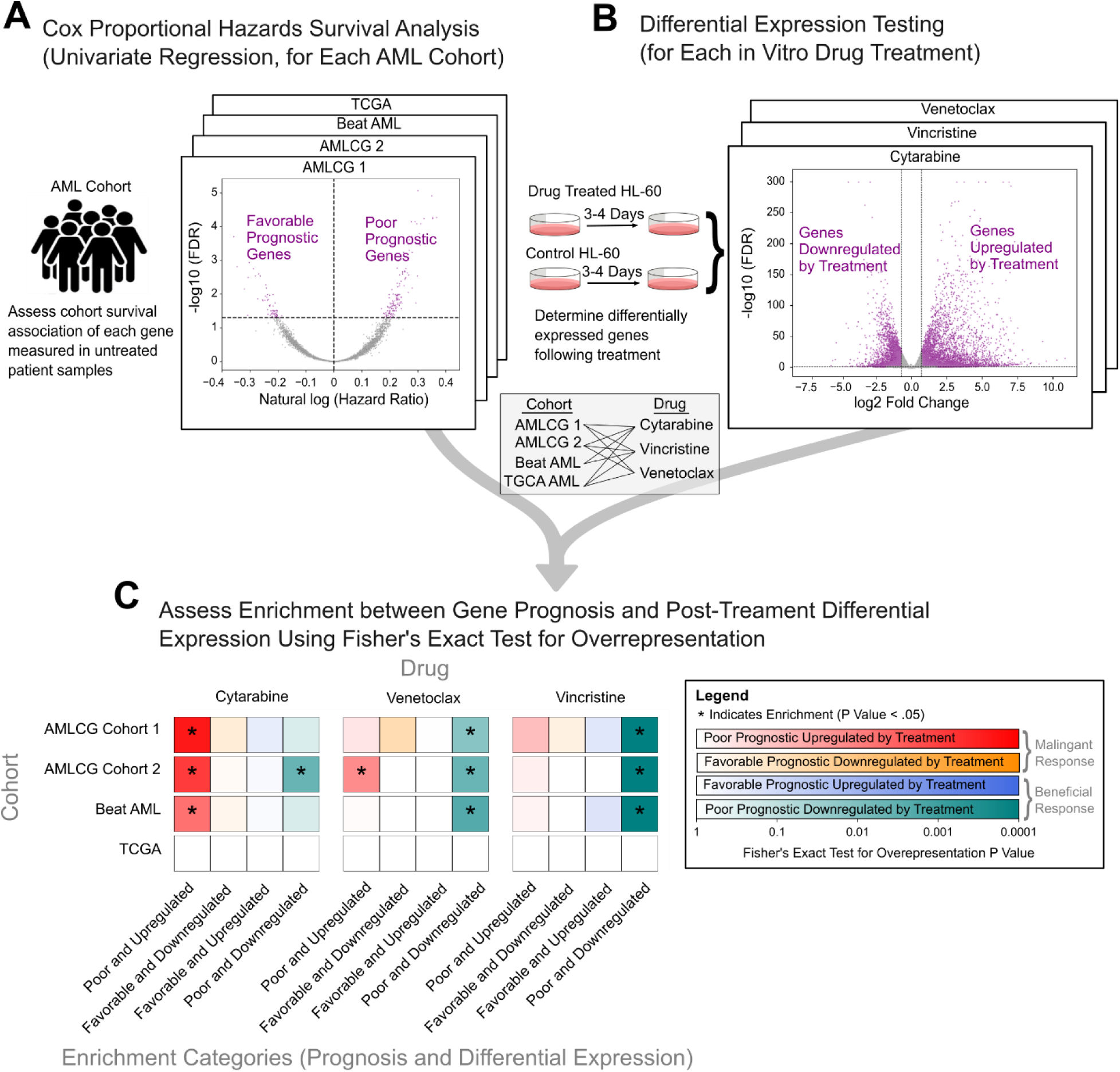
Statistical Enrichment Between Drug Induced and Prognosis Associated Genes. **A)** Genes measured in bulk RNA-seq of untreated leukemia patient samples were individually queried for prognostic value by assessing the association between gene expression levels and survival times using Cox proportional hazards regression (univariate), done separately in each of 4 cohorts: Beat, TCGA, AMLCG1 and AMLCG2. Shown are the results for AMLCG1 in the form of a volcano plot, with genes plotted according to natural log of hazard ratio (x-axis) and FDR (y-axis). Poor prognosis associated genes (natural log of hazard ratio > 0 & FDR < 0.05) and favorable prognosis genes (natural log of hazard ratio > 0 & FDR < 0.05) are indicated in purple. **B)** Genes were assessed for differential expression in bulk RNA-seq of HL-60 cells surviving in vitro chemotherapy treatment relative to untreated cells. This was done for 3 different drugs: cytarabine, venetoclax and vincristine. Shown are the results for cytarabine in the form of a volcano plot, with genes plotted according to log2FC (x-axis) and FDR (y-axis). Upregulated genes (log2FC > 0.7 & FDR < 0.05) and downregulated genes (log2FC > 0.7 & FDR < 0.05) are indicated in purple. **C)** Enrichment between prognostic association (poor or favorable) and differential expression (upregulated or downregulated) was assessed for each of 12 pairwise drug and cohort combinations, with drug specific differential expression and cohort specific prognosis associations assigned to each gene measured in both datasets. For each of 12 pairwise combinations (horizontal bars), Fisher’s exact test for overrepresentation was used 4 times (colored squares) to test for statistical enrichment (p < 0.05), once for each possible categorical overlap: upregulated and poor prognostic (red), downregulated and favorable prognostic (orange), upregulated and favorable prognostic (blue), downregulated and poor prognostic (green). Higher color intensity indicates a lower p value returned from Fisher’s exact test for overrepresentation and enrichment (p < .05) is denoted by an asterisk.

Significant enrichment (p value < 0.05) was observed in several overlap categories, with results most strongly driven by drug, rather than cohort choice (**Fig. 1C**) — specifically, cytarabine enrichment results were the most associated with malignant response in the form of poor prognostic genes being upregulated, while vincristine was predominantly associated with beneficial response in the form of poor prognostic genes being downregulated; venetoclax showed an intermediate association, with simultaneous (but weaker) enrichment for both of the above intersections. Enrichment of genes associated with favorable prognosis were less common, but downregulation was noted in the case of venetoclax in one cohort and upregulation in vincristine for another cohort.

Sensitivity analysis (**Supp. Fig. 6**) assessed the stability of these results: Fisher’s exact tests for overrepresentation were done using univariate (gene only) or multivariate (gene and available clinical covariates) survival regression, and FDR maximums for genes to be considered differentially expressed or prognosis-associated were varied between 0.01 and 0.1 (as 0.05 is the default in our above analyses). We found that multivariate regression shifted the more “malignancy-associated” drugs (cytarabine and venetoclax) towards beneficial responses while dampening such associations in vincristine. Less stringent FDR cutoffs generally strengthened existing associations, while more stringency nullified enrichment findings. With both types of parameter changes we observed some instability in terms of whether a specific drug and cohort remained enriched; however, broader drug-specific malignancy associations and general trends remained consistent. The major consequence of sensitivity analysis (under more stringent FDRs and multivariate regression) was the blurring of whether venetoclax or cytarabine is more associated with malignant or beneficial responses, although both drugs were still more associated with malignancy than vincristine which remains distinct in the upregulation of genes associated with beneficial response.

The drug most associated with treatment-induced malignancy based on upregulation of poor prognostic genes in non-killed cells, cytarabine, is also the most abundantly administered drug in all patient cohorts (**Supp. Fig. 7**) because of its superior efficacy. Venetoclax, a target selective drug that prevents BCL2-mediated survival, is used occasionally, and vincristine, the drug least associated with treatment-induced malignancy here has proven not effective in the clinic and is not used in standard of care (41,42).

### Common Genes Upregulated by Treatment and Linked to Poor Outcome

Of particular interest is the group of genes that were significantly upregulated in two out of the three HL-60 drug treatments and were found to be associated with poor outcomes in any cohort. Increasing the requirement for drug overlap revealed genes that were concordant in their effect directionality towards both upregulation by treatment and predicting poor prognosis (barring any strict fold change or FDR cutoffs). Assessment of the broader biological trends of drug-induced malignancy revealed 25 genes of interest (**Fig. 2**). All genes associated with survival in any cohort and also differentially expressed in any drug treatment are presented in **Supp. Table 1**.

**Figure 2:**
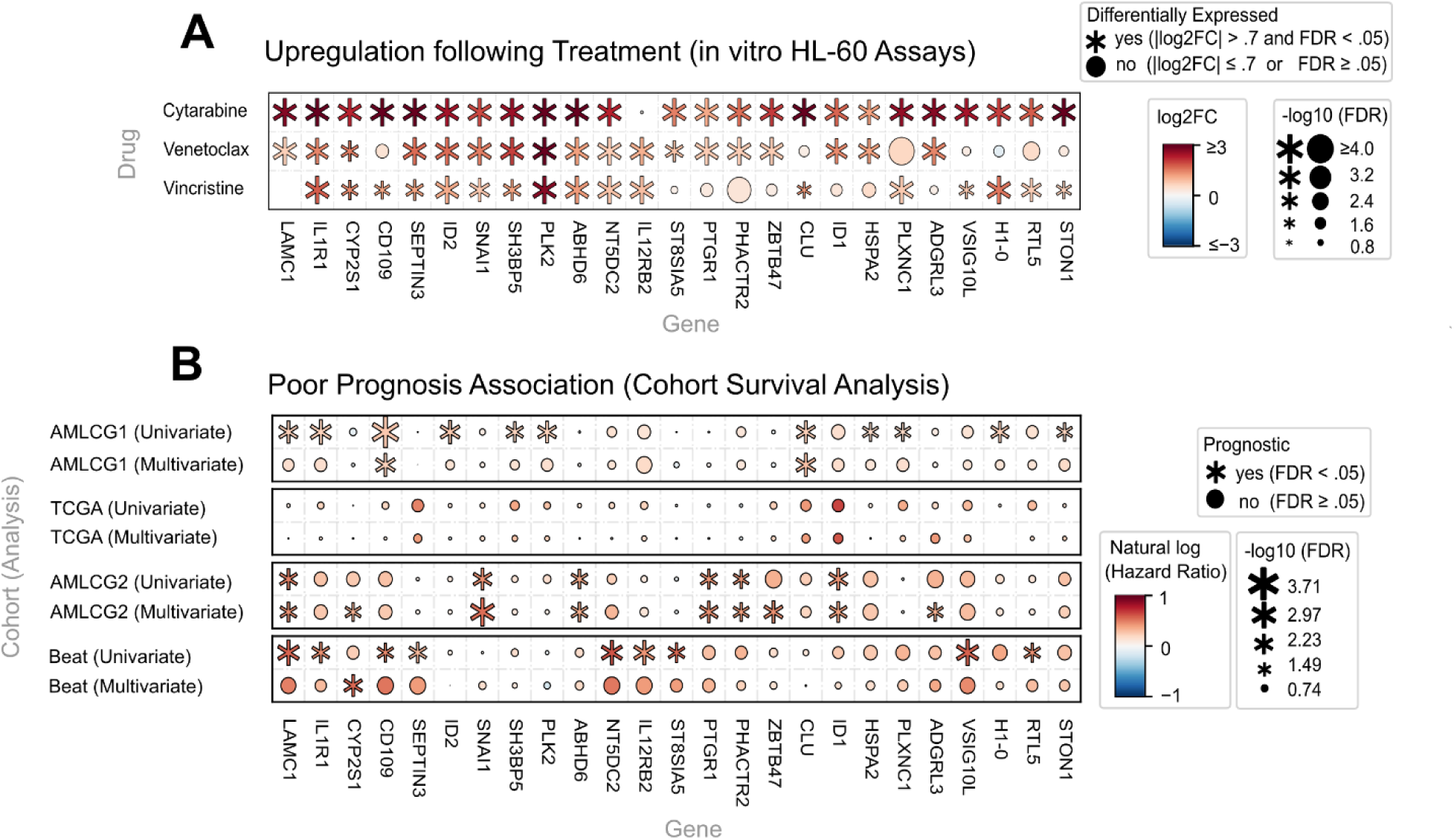
Poor Prognostic Genes Upregulated in the Majority of Chemotherapy Treatments. All genes shown are upregulated (log2FC > 0.7, FDR < 0.05) in at least 2/3 drugs and poor prognostic (natural log hazard ratio > 0, FDR < 0.05) in at least one cohort. **A)** Differential expression of genes (columns) in HL-60 surviving chemotherapy exposure, based on drug administered (rows). Gene expression fold change (log2) is indicated by color, with red indicating a positive fold change, blue indicating a negative fold change, and intensity indicating effect size. FDR is denoted by dot size, with larger dots indicating a lower FDR value. Genes considered differentially expressed (|log2FC| > 0.7 and FDR < 0.05) are marked with an asterisk, **B)** Prognostic association of genes (columns) in patient cohorts (rows). Covariates used in survival regression are further specified in row labels, univariate (gene only) or multivariate (gene, age, and white blood cell count if available). Prognostic association is indicated by color, with red indicating poor prognosis (natural log (hazard ratio) > 0), blue indicating favorable prognosis (natural log hazard ratio < 0), and intensity indicating effect size. A positive log hazard ratio indicates an increased death risk per unit gene expression and a negative log hazard ratio indicates the opposite, where gene expression is z- score normalized for comparability between genes. FDR is denoted by dot size, and prognosis associated genes (FDR < 0.05) are marked with an asterisk.

While few of these genes (*LAMC1*, *IL1R1*, *CYP2S1, CD109*), reached significant association with poor prognosis (natural log hazard ratio > 0, FDR < 0.05) in more than one cohort, almost all genes showed a proclivity towards poor survival (natural log of hazard ratio > 0, **Fig. 2B**). Similarly, of the genes significantly upregulated (log2FC > 0.7, FDR < 0.05) following at least two out of three drug treatments, only a portion were upregulated across all treatments (*IL1R1*, *CD109*, *SEPTIN3*, *ID2*, *SNAI1*, *SH3BP5*, *PLK2*, *ABHD6*, *NT5DC2*), but almost all genes showed a trend towards upregulation (log2FC > 0, **Fig. 2A**), thus motivating the generalization that these genes were upregulated by treatment and associated with poor prognosis.

### Transcription Factor Inference for Genes of Interest and Literature Searches

Next, we further evaluated the biological meaning of the genes that were exhibited upregulated by chemotherapy (in at least two out of three drugs) and were associated with poor prognosis (in any cohort).

Transcription factor (TF) inference using ChEA3 (**Fig. 3A**) shows a number of developmentally relevant, stress responsive and cancer-associated transcription factors (further discussed below) inferred to regulate of genes found to be upregulated by treatment and associated with poor outcome. In particular, the genes *CD109*, *IL1R1*, *ID1*, *ID2*, *PLK2*, *LAMC* and *CLU* seem to be jointly regulated by many of the top 20 transcription factors most highly inferred by CHEA3 (see methods). In the context of our drug treatments, this could indicate a concerted response. We note these TFs do not need to prognostic or differentially expressed themselves to regulate prognostic or differentially expressed genes returned by our analysis.

**Figure 3:**
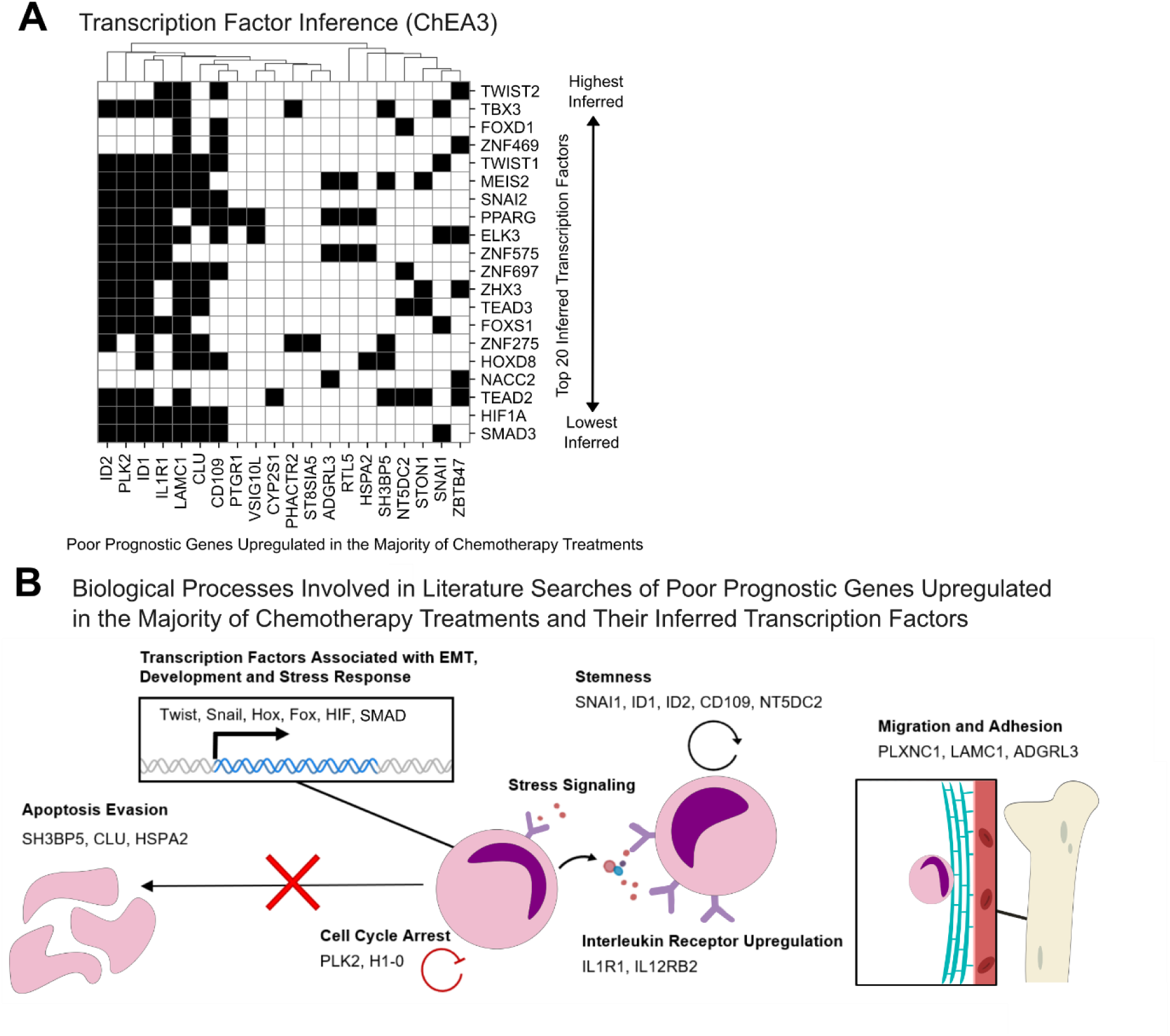
Transcription Factor Inference for Genes of Interest and Literature Searches. **A)** Transcription factor inference for poor prognostic genes (in any cohort) upregulated in the majority of chemotherapy treatments was done using the web based tool ChEA3, with interactions between inferred transcription factors and upregulated genes indicated by black squares on the clustermap. The top 20 inferred transcription factors are sorted by highest inference, decreasing along y-axis. The upregulated poor prognostic genes they regulate are hierarchically clustered along the x-axis to highlight common regulons. **B)** Graphical summary of literature associations between biological processes and poor prognostic genes (in any cohort) upregulated in the majority of chemotherapy treatments, plus their inferred transcription factors. Upregulated poor prognostic genes are associated with processes including stemness, cell cycle arrest, migration, adhesion, drug resistance, apoptosis evasion and stress signalling. Inferred transcription factors are commonly associated with EMT, development, and stress response.

Literature searches of genes upregulated in the treated cells and linked to poor outcome, predominantly in the context of AML but also in other cancers and hematological malignancies, suggested roles in the following biological processes: Stemness (*ID1*, *ID2*, *CD109*, *SNAI1*, *NT5DC2*), cell cycle arrest following stress response (*H1-0*, *PLK2*), apoptosis evasion following chemotherapy (*SH3BP5*, *CLU*, *HSPA2*), inflammation (*IL1R1*, *IL12RB2*), migration and adhesion (*LAMC1*, *ADGRL3*, *PLXNC1*), and retinoic acid metabolism (*CYP2S1*) relevant to differentiation blocks in acute promyelocytic leukemia and potentially other leukemias (18). The inferred upstream transcription factors were predominantly implicated in developmental processes (HOX, FOX, TEA), epithelial-to-mesenchymal transition (SNAIL, TWIST) and in stress response (*HIF1A*, *SMAD3*). These associations are depicted in **Fig. 3B**, and more comprehensively summarized in **Table 1** and **Table 2**, largely in the context of AML and hematopoietic malignancies where possible, but also of normal hematopoiesis and other cancers.

**Table 1:**
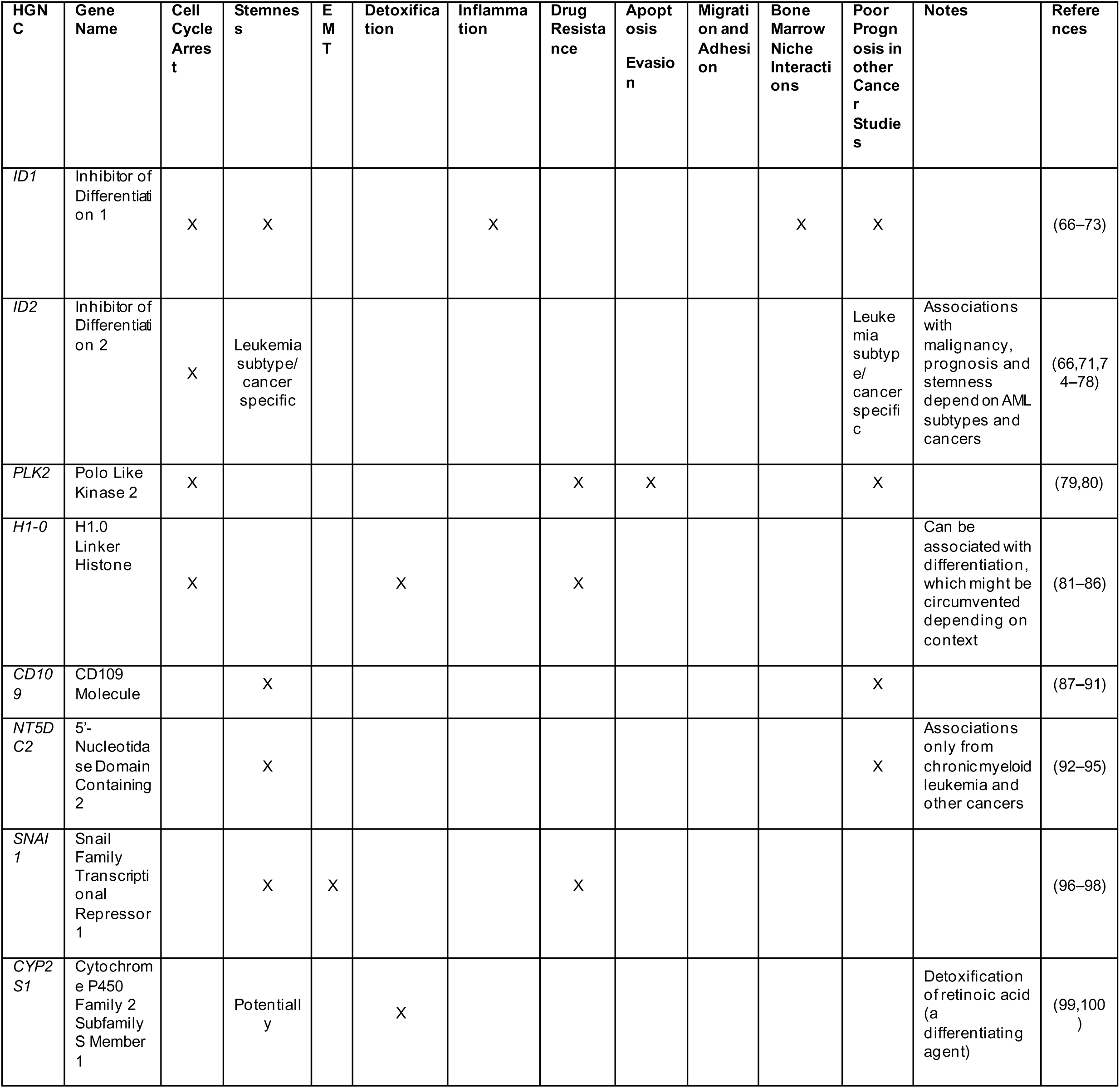
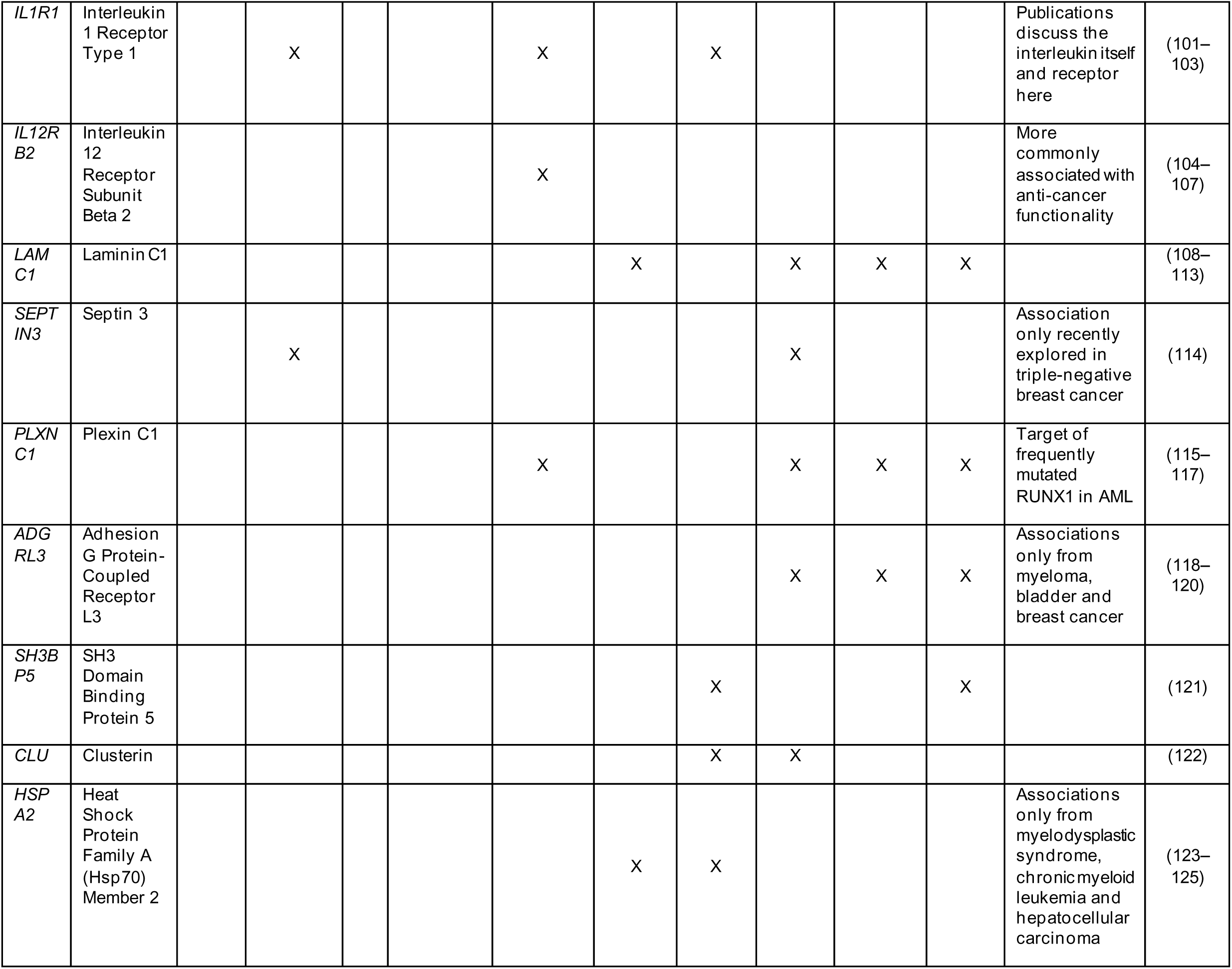
Biological Processes Associated with Poor Prognostic Genes Upregulated in the Majority of Drug Treatments (Fig. 2)

**Table 2:**
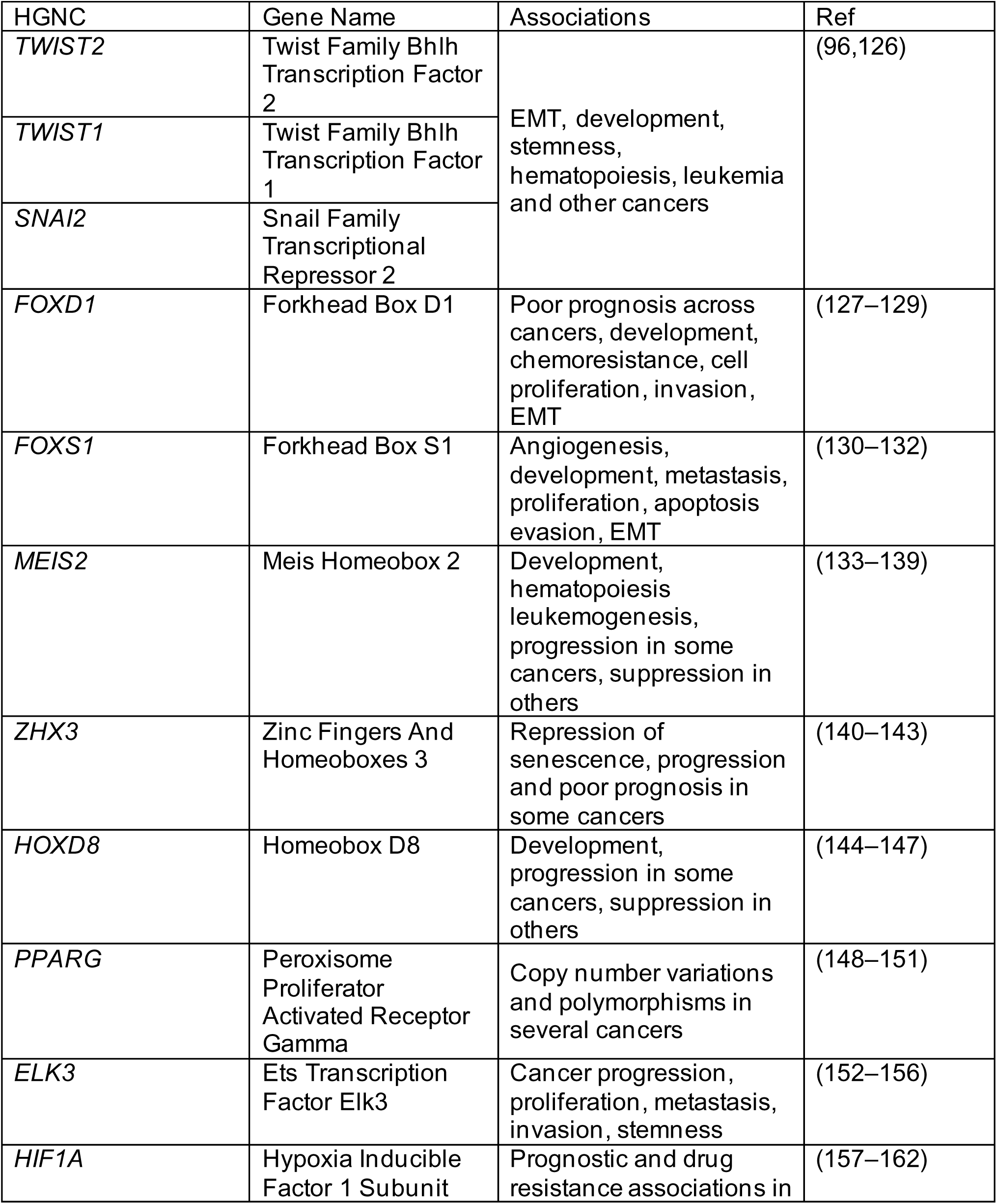
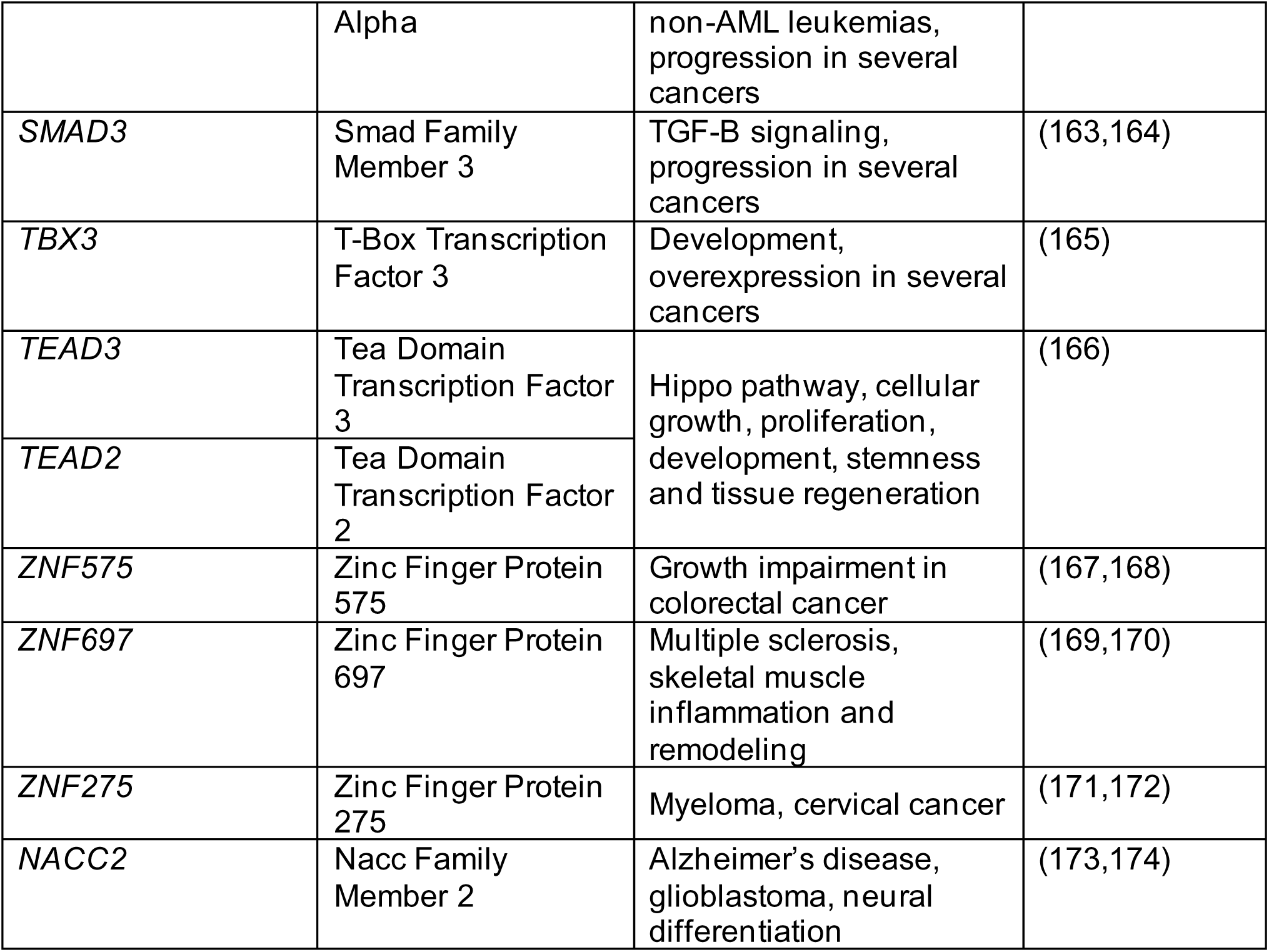
Top 20 Transcription Factors Inferred to Regulate Poor Prognostic Genes Upregulated in the Majority of Drug Treatments (. **Fig. 2)**

|  |  |  |  |
| --- | --- | --- | --- |
| HGNC | Gene Name | Associations | Ref |
| *TWIST2* | Twist Family Bhlh Transcription Factor 2 | EMT, development, stemness, hematopoiesis, leukemia and other cancers | (96,126) |
| *TWIST1* | Twist Family Bhlh Transcription Factor 1 |  |  |
| *SNAI2* | Snail Family Transcriptional Repressor 2 |  |  |
| *FOXD1* | Forkhead Box D1 | Poor prognosis across cancers, development, chemoresistance, cell proliferation, invasion, EMT | (127–129) |
| *FOXS1* | Forkhead Box S1 | Angiogenesis, development, metastasis, proliferation, apoptosis evasion, EMT | (130–132) |
| *MEIS2* | Meis Homeobox 2 | Development, hematopoiesis, leukemogenesis, progression in some cancers, suppression in others | (133–139) |
| *ZHX3* | Zinc Fingers And Homeoboxes 3 | Repression of senescence, progression and poor prognosis in some cancers | (140–143) |
| *HOXD8* | Homeobox D8 | Development, progression in some cancers, suppression in others | (144–147) |
| *PPARG* | Peroxisome Proliferator Activated Receptor Gamma | Copy number variations and polymorphisms in several cancers | (148–151) |
| *ELK3* | Ets Transcription Factor Elk3 | Cancer progression, proliferation, metastasis, invasion, stemness | (152–156) |
| *HIF1A* | Hypoxia Inducible Factor 1 Subunit | Prognostic and drug resistance associations in | (157–162) |
|  | Alpha | non-AML leukemias,<br>progression in several<br>cancers |  |
| *SMAD3* | Smad Family<br>Member 3 | TGF-B signaling,<br>progression in several<br>cancers | (163,164) |
| *TBX3* | T-Box Transcription<br>Factor 3 | Development,<br>overexpression in several<br>cancers | (165) |
| *TEAD3* | Tea Domain<br>Transcription Factor<br>3 | Hippo pathway, cellular<br>growth, proliferation,<br>development, stemness<br>and tissue regeneration | (166) |
| *TEAD2* | Tea Domain<br>Transcription Factor<br>2 |  |  |
| *ZNF575* | Zinc Finger Protein<br>575 | Growth impairment in<br>colorectal cancer | (167,168) |
| *ZNF697* | Zinc Finger Protein<br>697 | Multiple sclerosis,<br>skeletal muscle<br>inflammation and<br>remodeling | (169,170) |
| *ZNF275* | Zinc Finger Protein<br>275 | Myeloma, cervical cancer | (171,172) |
| *NACC2* | Nacc Family<br>Member 2 | Alzheimer's disease,<br>glioblastoma, neural<br>differentiation | (173,174) |

### Single-Cell RNA-seq of HL-60 Response to Chemotherapy Treatment

The dichotomous response of tumor cell populations, with the intended killing of some cells and unintended induction of genes that may promote recurrence in other cells underscore the importance of cell phenotype heterogeneity within (possibly isogenic) cancer cell populations. Thus, we next sought to examine the transcriptome changes in the HL-60 cells at single-cell resolution using single-cell RNA-seq following chemotherapy with either vincristine (5 nM, greater than EC50 concentration) or cytarabine (1.6 μM, EC50 concentration). These two drugs and doses were explored in two separate experiments (“Experiment A” for vincristine and cytarabine, and “Experiment B” for vincristine alone).

Following 48h of treatment with either vincristine or cytarabine (Experiment A), we observe a global shift in transcriptome state between untreated (blue, **Fig. 4A**) and treated cells (green: vincristine, red: cytarabine). The transcriptome shift was most pronounced in cytarabine treatment where treated and untreated populations were almost mutually exclusive, whereas separation was less pronounced after vincristine treatment, as a substantial portion of cells remained in transcriptome states characteristic of untreated cells. Additionally, we observe a few untreated cells already occupying clusters typical of treated cells in both treatments.

**Figure 4:**
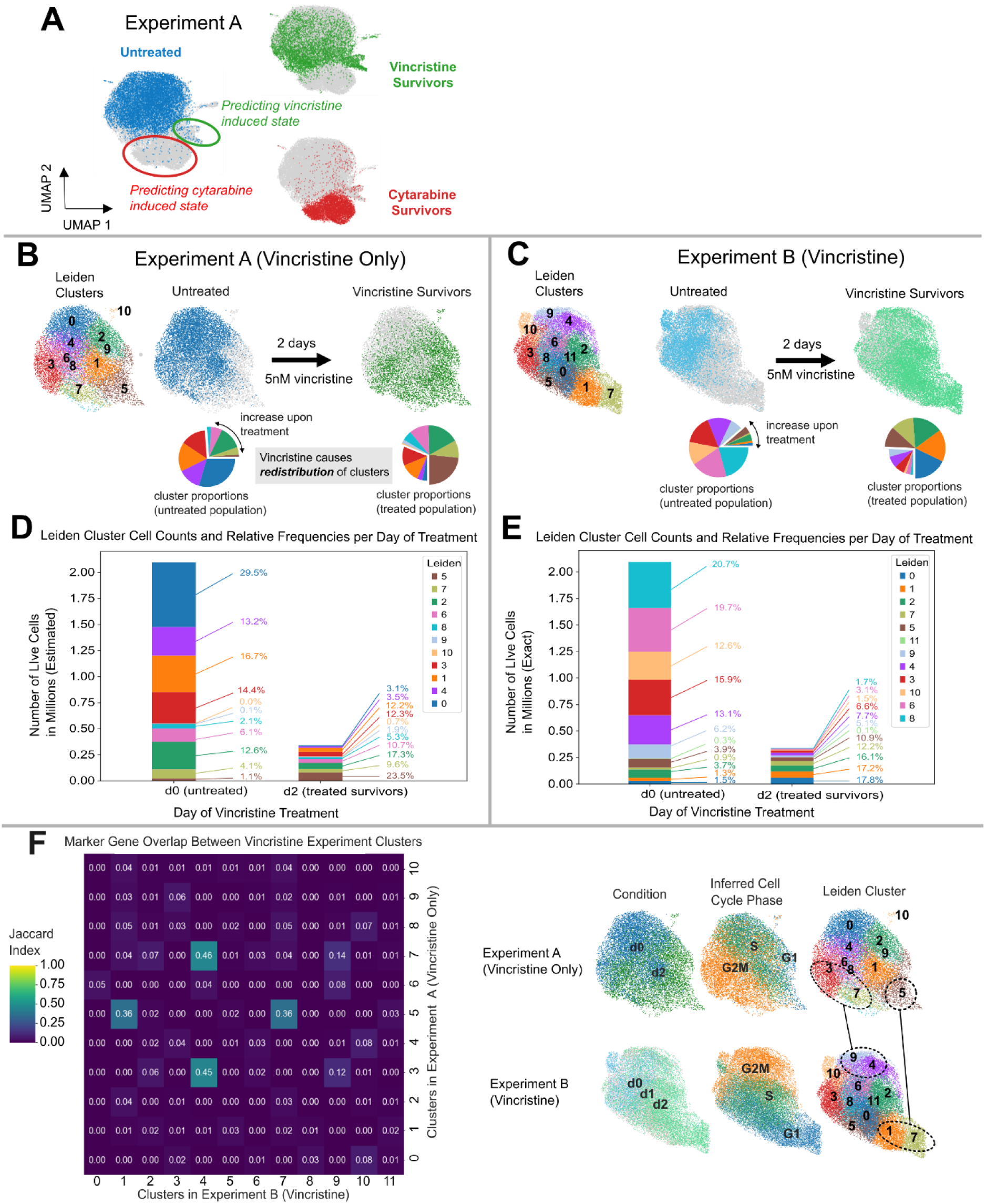
Single-Cell RNA-seq of HL6O Response to Chemotherapy Treatment. **A)** Single-cell RNA-seq of HL-60 Surviving 48 hours of chemotherapy with cytarabine or vincristine (both treatments done in parallel in Experiment A) reveals shifts in treated population towards transcriptomic States that were previously occupied by only a small fraction of untreated cells. Cell transcriptomes (dots, grey by default) are pooled in UMAP space, with untreated cells in blue, cytarabine treated cells in red and vincristine treated cells in green. **B,C)** Untreated HL-60 and vincristine treated survivors in two separate single-cell RNA-seq experiments. Leiden clustering of cells in each experiment (assignments shown in UMAP space) and calculation of cluster proportions in cell populations before and after treatment reveals a shift in cluster occupancy distribution, indicated by pie charts. Specifically, an increase in the proportions of clusters uncommon prior to treatment (offset and expanded wedges) and corresponding decrease in clusters common in the untreated populations. **D,E)** Occupancy (in live cell numbers) of Leiden clusters generated for vincristine treated survivors and untreated HL-60 in Experiment A and Experiment B, respectively. Stacked bar charts indicate absolute number of cells before and after vincristine treatment (bar height), and percent occupancies of clusters within a given condition are labelled. Several clusters decrease in cell number and proportion of cell population following treatment, while others increase cell in number and proportion. Some clusters increase in proportion following treatment despite a decrease in absolute number. Live cell counts before and after treatment were available for Experiment B, and used to estimate cell counts in Experiment A. **F)** Cluster marker gene overlap between all cell clusters in Experiment A and Experiment B vincristine treatments, assessed using the Jaccard index for each comparison. Marker genes are assigned by testing differential expression in each cluster against all others within an experiment (Welch’s t-test): the top 100 markers (log2FC > 0.7 and FDR value < 0.05) were used, sorted by increasing FDR value. The heatmap shows overlap coefficients between clusters. Circled and connected UMAP clusters are those showing marker gene correspondence (Jaccard > .1) between experiments.

Next, we focus exclusively on the vincristine treatment data of Experiments A and B (**Fig. 4 B, C,** respectively). To determine whether a significant population - wide cell state shift occurred with vincristine treatment, we quantified the relative occupancy of treated and untreated cells within each of the individual Leiden clusters (cells with similar transcriptomes), identified by standard single-cell transcriptome cluster analysis (see methods). In both experiments, clusters common in the untreated culture decreased in population proportion following vincristine treatment, with a corresponding increase of cells in the clusters less common in the untreated culture. Thus, the change induced by treatment can be characterized as a redistribution of relative cluster occupancy, rather than a de novo generation of new cell states (clusters).

From single-cell RNA-seq alone, we can only infer the relative proportions of clusters in a sample. To obtain absolute cell numbers in each cluster, cluster proportions were scaled by live cell counts recorded for each sample (live cell counts were only recorded for Experiment B and used as a proxy for Experiment A). This allowed us to obtain insight into the absolute population dynamics following drug treatment (**Fig. 4 D, E**), revealing several clusters that increase in cell number despite most clusters shrinking from cell death.

While batch effects (from sample preparation) notoriously make the combining of cells and replication of clusters between independent experiments challenging, we could establish correspondence by determining marker genes for all clusters, and calculating the Jaccard index to compare cluster marker genes across experiments (**Fig. 4F**). Marker genes for a cluster were defined as the top 100 genes sorted by increasing FDR when testing a given cluster against all other clusters in a given experiment (provided log2FC > 0.7 and FDR < 0.05). Clusters most correspondent with one another across experiments were those associated with (i) G2 or mitosis, (ii) the clusters most populated by treated cells and least populated by untreated cells. Overlapping marker genes between clusters are provided in **Supp. Table 2**.

Taken together, these changes reflect widespread chemotherapy induced shifts in surviving cells towards phenotypes that pre-existed before the treatment but were latent, that is, uncommon in the untreated population. This result revealed the immediate, diverging response of a heterogeneous cell population as a combination of distinct cell state transitions induced by treatment, but did not determine which initial state contributed most to a given transition given that phenotypic heterogeneity underlies differential sensitivity to perturbations (2,43–45), nor how the new population distribution contributes to longer-term change of the biology of the populations, such as treatment resistance. Addressing these questions would require physical isolation and re-culturing of the individual clusters (46).

## Discussion

We examined transcriptome changes in HL-60 leukemic cancer cells that survived chemotherapy treatment for molecular signs of cancer promoting responses in view of the burgeoning notion that cancer treatment is a double-edged-sword: While current treatment kills tumor cells, hence shrinking the tumor in the short term (which may address an acute clinical need), such cytocidal treatment also harbors an element of disease stimulation: it may increase relapse risk if the surviving cells are converted to a more malignant phenotype in response to cell stress and tissue injury inflicted by the treatment. The study was not designed to expose and quantify the contribution of selection of rare pre-existing variant cells caused by random mutations or epigenetic alterations that confer resistance to treatment – the classical explanation for recurrence of treatment-resistant cancer after initial remission. Instead, our goal was to examine the rapid “mass” shift (i.e. of many individual cells at a cell population scale) of cell phenotype and to study the biological significance of the drug-induced cellular programs in the residual cancer cells.

Specifically, we sought to determine if the rapid upregulation of genes that takes place in a large fraction of the surviving cell population after treatment may offer an explanation for the double-edged effect of cancer treatment (6,47). Thus, going beyond the conventional identification of differentially expressed genes (DEG) by comparing treated and untreated cells, we asked if among upregulated DEG there were instances of genes associated with poor prognosis in leukemia patients.

Unlike other studies describing the phenomenon of treatment-promoted progression, we went beyond searching for gene signatures among the DEG that indicated higher malignancy in cells that have survived treatment. Instead, to avoid bias in gene function annotations, we asked if the genes significantly upregulated after treatment with three canonical drugs in the non-killed leukemic cells were also significantly associated with shorter survival times when found to be highly expressed in treatment-naïve samples of patients diagnosed with the same cancer given that such transcriptome data is widely available (in contrast to those from post-treatment samples).

The robust upregulation by three drugs in AML cells of dozens of genes associated with poor prognosis in AML cohorts (**Fig. 2**) is consistent with the notion that cancer therapy may inherently be a trade-off between tumor suppression due to the intended cytocidal effects and the unintended treatment-induced progression towards higher malignancy in the non-killed but injured cells. Chemotherapy can achieve lasting remission if the therapeutically intended eradication of a large number of cells outweighs the unintended increase in cancer cell malignancy in a small number of surviving cells. In the transcriptomes of cells surviving chemotherapy, we also find instances of differential expression of genes that may contribute to beneficial patient outcomes, namely poor prognostic genes being downregulated or favorable prognostic genes being upregulated (**Supp. Table 1**).

Literature searches and transcription factor inference further support the idea that many genes upregulated by chemotherapy in our cells may indeed contribute to treatment-induced increase in aggressiveness, in view of their known roles stemness, EMT, inflammation, apoptosis evasion, stress-response, etc. (**Fig. 3B**, **Table 1**, **Table 2**).

Going beyond traditional gene function annotation-based analysis, our study design is centered around the intersection of two statistical associations: genes associated with treatment response in vitro and survival in clinical cohorts; while avoiding the bias from biological interpretation of gene function, this scheme calls for caution in interpretation because of experimental and statistical limitations.

First, the RNA-seq experiments were designed with a short exposure of 2-4 days at doses above the half-maximal effective concentrations for cell death (EC50). This time frame is short relative to what would be required for selection of pre-existing (genetic), or even of induced cell phenotypes, that would enrich for cells with more aggressive phenotypes. Thus, the fundamental distinction between active transcriptomic response due to cell intrinsic regulation and passive change of abundance of particular cell phenotypes due to population dynamics was not explored here (2,16,21,22,24) — although we cannot exclude contributions by the latter towards treatment-associated progression at longer time scales. Similarly, our limited exploration of drug doses does not provide insight into the non-monotonical dose-responses influencing the above-mentioned population dynamics (48). As discussed elsewhere, detailed quantitative analyses and mathematical models are needed to distinguish between regulated induction or selection for a more malignant phenotype (2,48–52).

In conjunction with the short-time scale, the cells surviving treatment in our experiment were not sorted and assayed for increased resistance or even malignancy such as clonogenicity to demonstrate an effective increase in malignancy. Another functional aspect not studied is paracrine signaling emanating from non - resistant dying cells and debris following their death; such “danger signals” are thought to trigger adversarial tumor responses (53–57) and may have contributed to the shift in gene expression programs towards malignancy.

A second caveat in the interpretation pertains to the patient cohort survival analysis that we used to assign a prognostic meaning to the genes identified as DEG between treated and untreated cells in vitro. We took this approach to avoid relying on the often - biased gene function annotations and literature (selection bias) to ascertain contributions of the upregulated genes to increased malignancy. Instead, malignancy was derived directly from associations between patient gene expression at diagnosis and survival time.

The central assumption in this analysis was that the cell populations pushed towards a state of higher malignancy by treatment-stress would in some fashion resemble tumor features at diagnosis that indicate a worse survival outcome. As with any cohort study that seeks to predict prognosis at the level of individual genes at baseline, this comes with its own caveats. The more specific postulate is that the genes expressed at higher levels in treatment-naïve AML cells have the same detrimental impact on survival as those genes that we observed in our in vitro experiments to be upregulated by treatment (and vice versa). While this assumption may be plausible for some general properties of oncogene or tumor-suppressors, as it is widely, albeit tacitly, held in cancer genetics, it may not hold for specific biological functions embodied by the specific DEG in our respective clinical association studies. The concluding analysis of gene annotations and literature searches, which we saved only for this final step however, lends support to our hypothesis: We found that genes upregulated in HL-60 cells in vitro by treatment and that were also associated with poor prognosis when highly expressed in the pretreatment AML patients have known roles in malignant processes (**Fig. 3**, **Table 1**, **Table 2**). This result would be consistent with the concept of treatment-induced progression, thus corroborating the broader idea that cytocidal treatment can backfire.

Thirdly, as to the choice of drugs, it was intriguing to see how robustly the specifics of the cytocidal intervention influenced the gene enrichment results. The drug most associated with treatment-induced malignancy was cytarabine, which shifted gene expression toward malignancy associated genes by an overwhelming margin, followed by venetoclax, and least, vincristine. Cytarabine, the most commonly administered drug among these three, is the backbone of standard of care “7+3” treatment in AML (58,59). It is conceivable that different drugs with different mechanisms of action in killing could shift surviving cell populations differentially towards more alternative malignant states (**Supp. Fig. 1**). Of note is the much more muted response in treatment-induced malignancy for vincristine: This drug is widely studied for resistance development in AML as it induces the detoxification system, including upregulation of MDR1 (also known as *ABCB1*) efflux pump that ejects the drug (2). In doing so, vincristine may possibly reduce cell stress, but it also has minimal clinical efficacy (42). Thus, it is may be telling that our experiments with three drugs revealed that the gradient of clinical efficacy, at least in inducing initial remission (cytarabine > venetoclax > vincristine), scales with the extent of cell state shifts that these drugs trigger in the surviving cells, underscoring the inescapable intrinsic constraints of the double-edged sword effect: More efficient killing is associated with more cell stress in the surviving cells.

Conversely, the choice of cohort seemed to not influence the gene enrichment results as strongly, despite minimal overlaps of the prognostic genes observed between cohorts (**Supp. Figs. 2, 3**). One explanation for this disparity could be cohort-specific biological differences that were not accounted for (patient sampling, disease heterogeneity, varying drug treatment after the initial gene expression measurement that we study). Low to absent overlap of prognostic marker signatures in oncology is all too often observed (60). Nevertheless, here the distinct genes identified to be associated with outcome in each cohort was informative enough to capture more general associations of either malignant or beneficial leukemic response when assessed alongside our drug treatment data.

Finally, in addition to the cautionary interpretation entailed by our study design, there are methodological limitations with the enrichment analysis. While the outcomes of the contingency tests reflect the general trend towards or away from a malignancy-increasing response in the treated leukemic blasts, simplifying assumptions and limitations are worth noting. First, differential expression and association with survival are assessed on a gene-by-gene basis, treating genes independently of each other when in reality, genes act in concert due to regulatory interdependence. Second, when the contingency table was constructed, we treated the results of differential expression and survival analysis as “ground truth”, set by fold-change and FDR cutoffs, yet these inferences could be incorrect (type 1 errors can at least be controlled, but type 2 errors are uncontrolled). This makes detecting a true dependence in the contingency table more difficult and could be accounted for in the future using a more complex inference model (61). Third, enrichment results only indicate if the number of items in a given category is greater than expected assuming independence between the categorical variables; they do not provide insight into the biological importance of different genes in our test, or the relative importance of different contingency table intersections. For instance, we might observe a lack of enrichment for genes associated with poor prognosis with the genes upregulated in our experiments, suggesting that the experimentally observed upregulation by a drug does not contribute to malignancy. However, it might counterintuitively be the case that such upregulated poor prognostic genes were particularly malignant, sufficient to drive resistance and relapse, yet present only at a very low number in the category of “significantly upregulated”, as might be the case with a handful of stemness related transcription factors.

A final limitation of the statistical analysis pertains to the perennial challenges with cohort survival association, including but not limited to confounders, mediators, decoupling of phenotype from survival time, evolving treatment regimens and patient heterogeneity. Even where covariates are accounted for, overfitting can occur (62–65).

In conclusion, despite the methodological limitations, this study offers substantial molecular support of the emerging idea that treatment intended to reduce tumor burden by killing cancer cells, can, due to the inevitable presence of surviving cells stressed by the treatment, undergo cell phenotype changes that confer higher malignancy. Hence, these cells may contribute a treatment-induced component to progression and relapse.

Our findings are of great significance for drug discovery: In the quest for more potent and selective cancer drugs, we need to go beyond screening for efficacy of killing cancer cells and consider the qualitative nature of phenotype alterations in surviving cells effected by the new drug. Considering unintended biological responses may pave the way to minimizing the double-edge sword effect immanent to cytocidal therapy – an untapped room for improvement of therapeutic efficacy that is not simply doubling-down on cell killing.

## Acknowledgements

INSERT FUNDING INFORMATION HERE

---

**Supplementary Figure 1:**
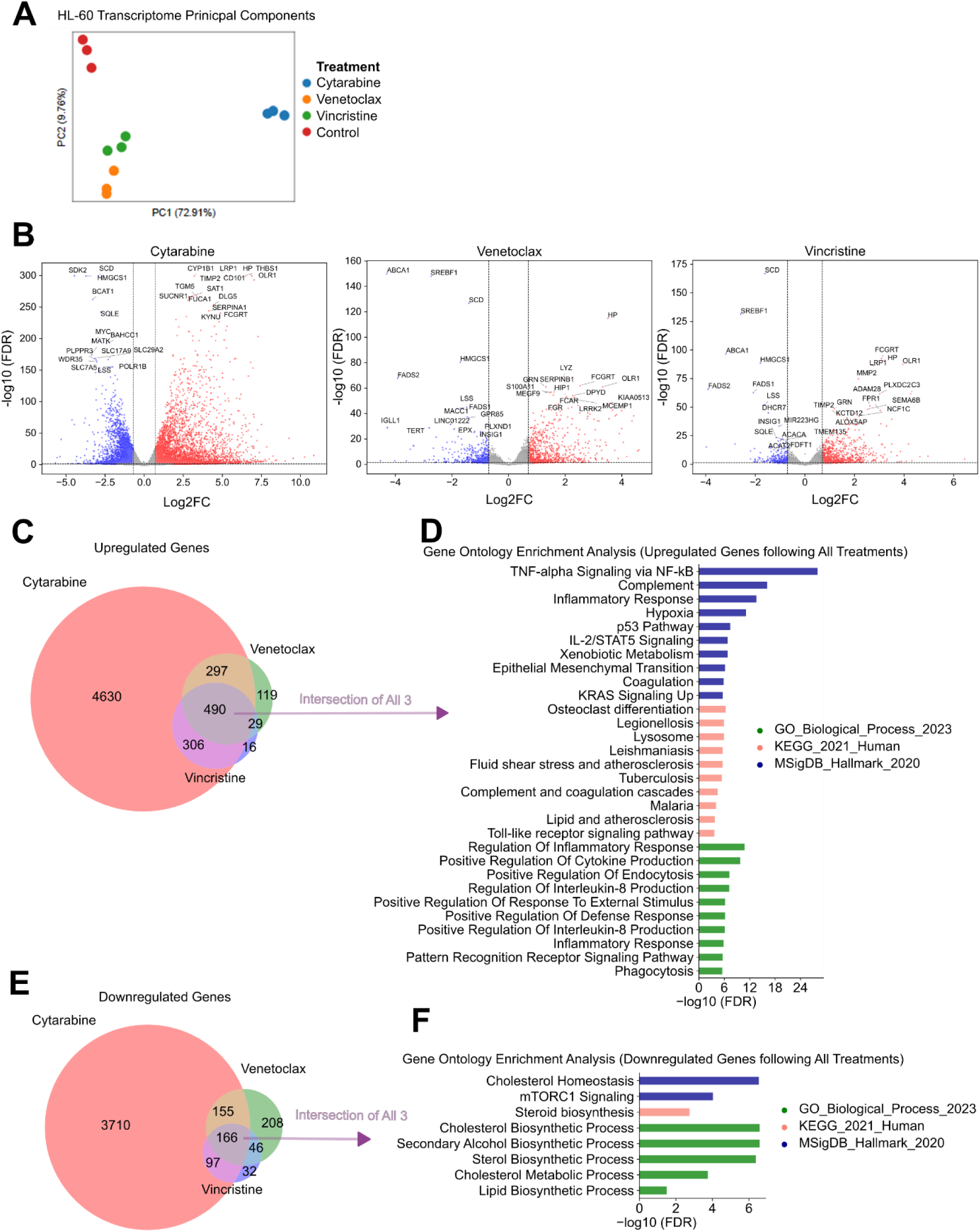
Transcriptome Changes in HL-60 Following Chemotherapy Treatment. **A)** Bulk-seq transcriptomes of HL-60 surviving chemotherapy treatment (cytarabine, vincristine or venetoclax) and untreated control, shown along first 2 principal components with variance explained for each component indicated as percentage. 3 biological replicates per condition. **B)** Volcano plots of differentially expressed genes following each of 3 chemotherapy treatments. Upregulated genes (log2FC > 0.7, FDR < 0.05) are shown in red, downregulated genes (log2FC < -0.7, FDR < 0.05) in blue. Top 15 genes (lowest FDR value) for each category are annotated. **C)** Venn diagram showing overlap sizes and overall numbers of genes upregulated in different treatments. **D)** Gene ontology enrichment for upregulated genes in all treatments (intersection) and corresponding processes in select pathway/gene set databases (top 10 results per database, FDR < 0.05) **E, F)** Identical to C, D respectively, only with downregulated genes assessed instead of upregulated genes.

**Supplementary Figure 2:**
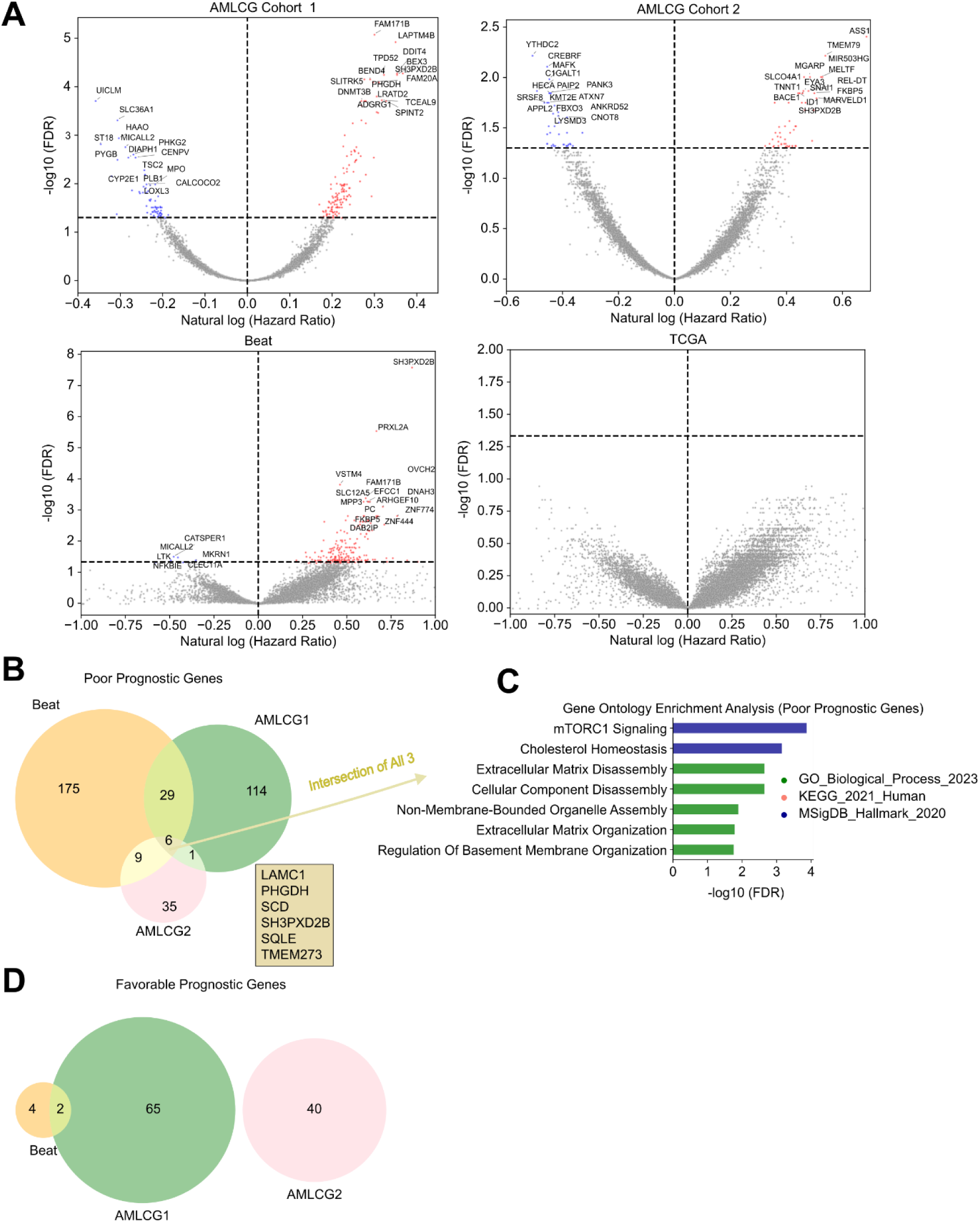
Cohort Survival Analysis (Univariate) **A)** Volcano plots of prognosis associated genes in each of 4 acute myeloid leukemia cohorts after univariate Cox regression (each gene is fitted as the sole covariate, and FDR corrected). Poor prognosis associated genes (natural log hazard ratio > 0, FDR < 0.05) are shown in red, favorable prognosis associated genes (natural log hazard ratio < 0, FDR < 0.05) are shown in blue. Top 15 genes (lowest FDR value) for each category are annotated. **B)** Venn diagram showing overlap sizes and overall numbers of poor prognosis associated genes in each cohort. TCGA returned no results. **C)** Gene ontology enrichment for poor prognosis associated genes in all 3 remaining cohorts (intersection) and corresponding processes in select pathway/gene set databases (top 10 results per database, FDR < 0.05) **D)** Venn diagram showing overlap sizes and overall numbers of favorable prognosis associated genes in each cohort. TCGA returned no results.

**Supplementary Figure 3:**
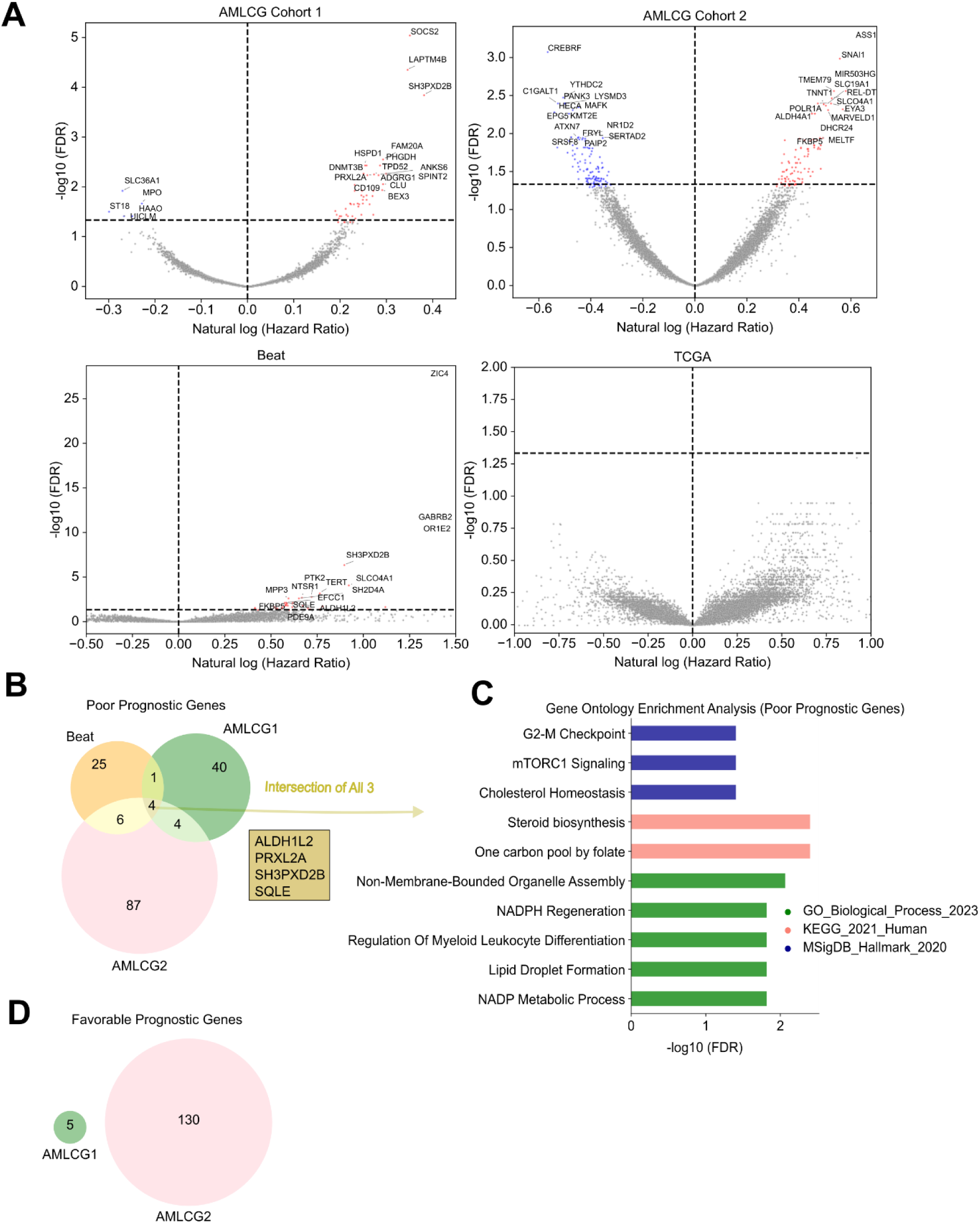
Cohort Survival Analysis (Multivariate) **A)** Volcano plots of prognosis associated genes in each of 4 acute myeloid leukemia cohorts after multivariate Cox regression (each gene is fitted as a covariate, along with age and white blood cell count, if available, and FDR corrected). Poor prognosis associated genes (natural log hazard ratio > 0, FDR < 0.05) are shown in red, favorable prognosis associated genes (natural log hazard ratio < 0, FDR < 0.05) are shown in blue. Top 15 genes (lowest FDR value) for each category are annotated. **B)** Venn diagram showing overlap sizes and overall numbers of poor prognosis associated genes in each cohort. TCGA returned no results. **C)** Gene ontology enrichment for poor prognosis associated genes in all 3 remaining cohorts (intersection) and corresponding processes in select pathway/gene set databases is shown to the right (top 10 results per database, FDR < 0.05) **D)** Venn diagram showing overlap sizes and overall numbers of favorable prognosis associated genes in each cohort. TCGA and Beat returned no results.

**Supplementary Figure 4:**
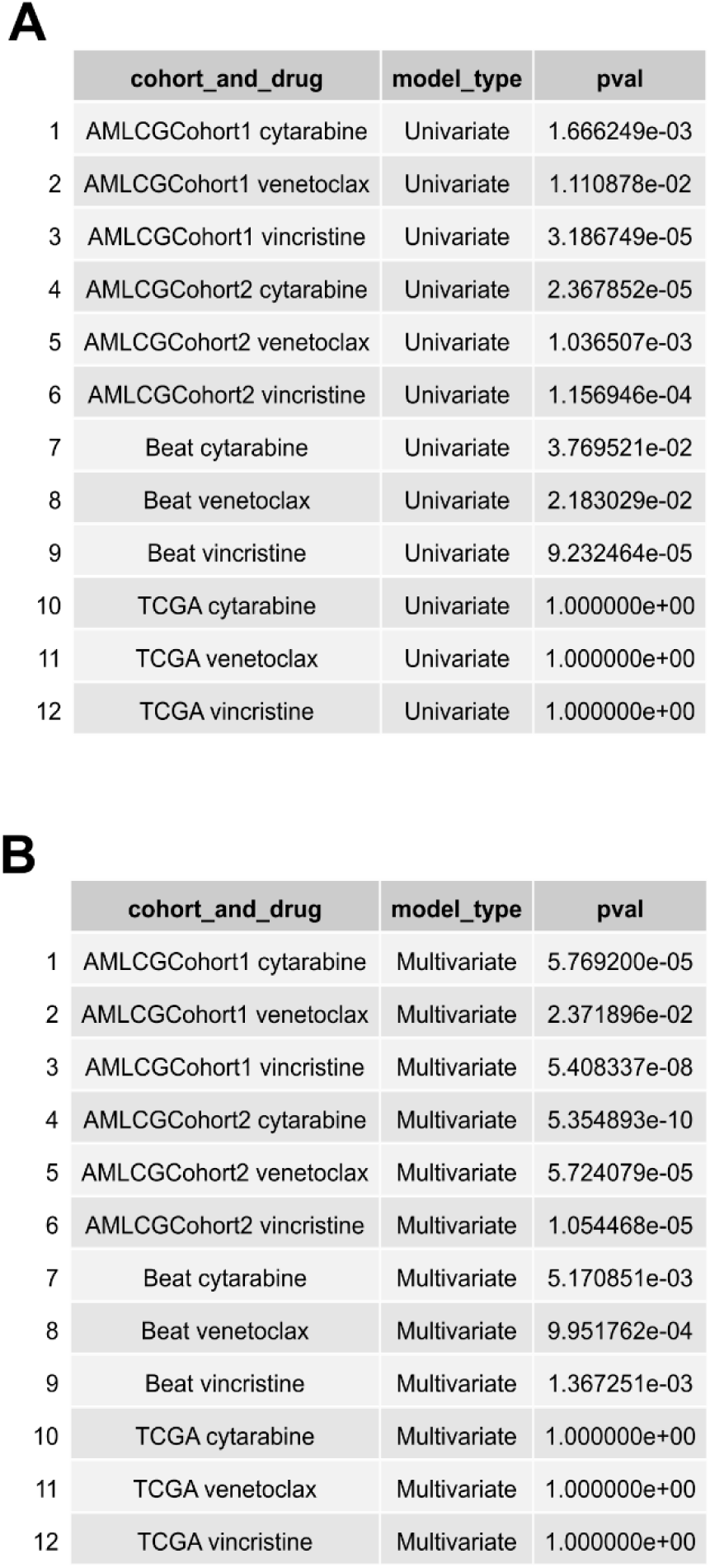
Enrichment between Gene Prognostic Association and Differential Expression Following Chemotherapy using Fisher’s Exact Test for 3x3 Contingency Tables. Low p values indicate a lack of statistical independence between gene prognostic association (cohort specific) and differential expression (drug specific), assessed using contingency tables for the categorizations of “oor”, “favorable”, or “no association” for prognosis, and “upregulated”, “downregulated” or “no differential expression” for differential expression. **A)** P values for each cohort and drug, where prognosis association was determined with individual genes as the sole covariates (univariate). **B)** P values for each cohort and drug, where prognosis association was determined with individual genes, patient age, and white blood cell count if available, as covariates (multivariate).

**Supplementary Figure 5:**
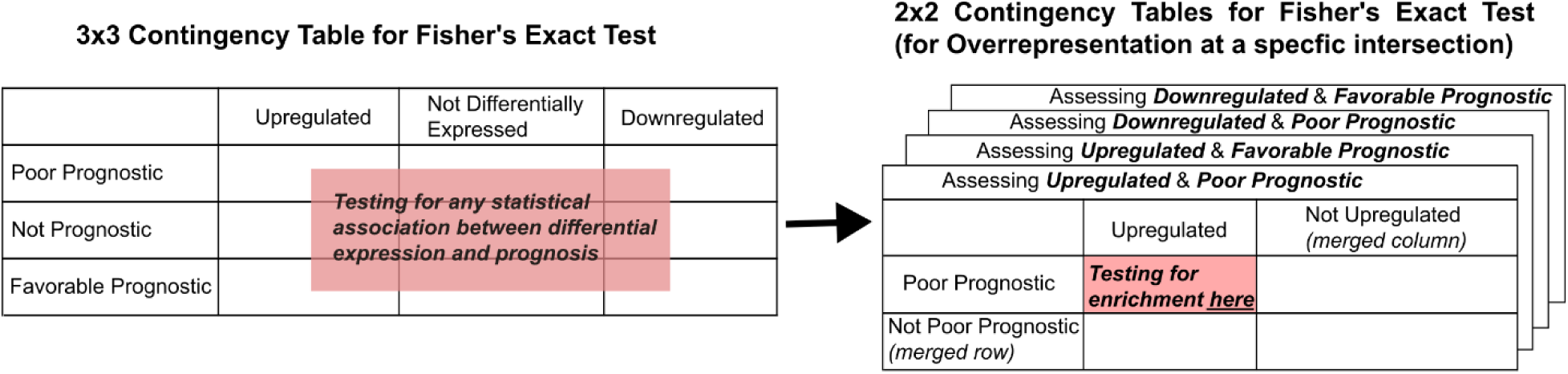
Collapsing 3x3 Contingency Tables to 2x2 for Fisher’s Exact Test (for Overrepresentation) Following initial Fisher’s Exact Tests for 3x3 contingency tables to test for any (unspecified) associations, contingency tables were collapsed into 2x2 tables to perform Fisher’s Exact Test for overpresentation, testing for specific associations between differentially expressed and prognosis associated genes. This was done by electing a differential expression category (column) and prognosis association category (row) of interest, then merging the remaining rows and columns, as illustrated for the example assessing the overlap between “upregulated” and “poor prognostic” genes. This was repeated 3 more times to test for additional enrichments (“upregulated and favorable prognostic”,“ downregulated and poor prognostic”, “downregulated and favorable prognostic”).

**Supplementary Figure 6:**
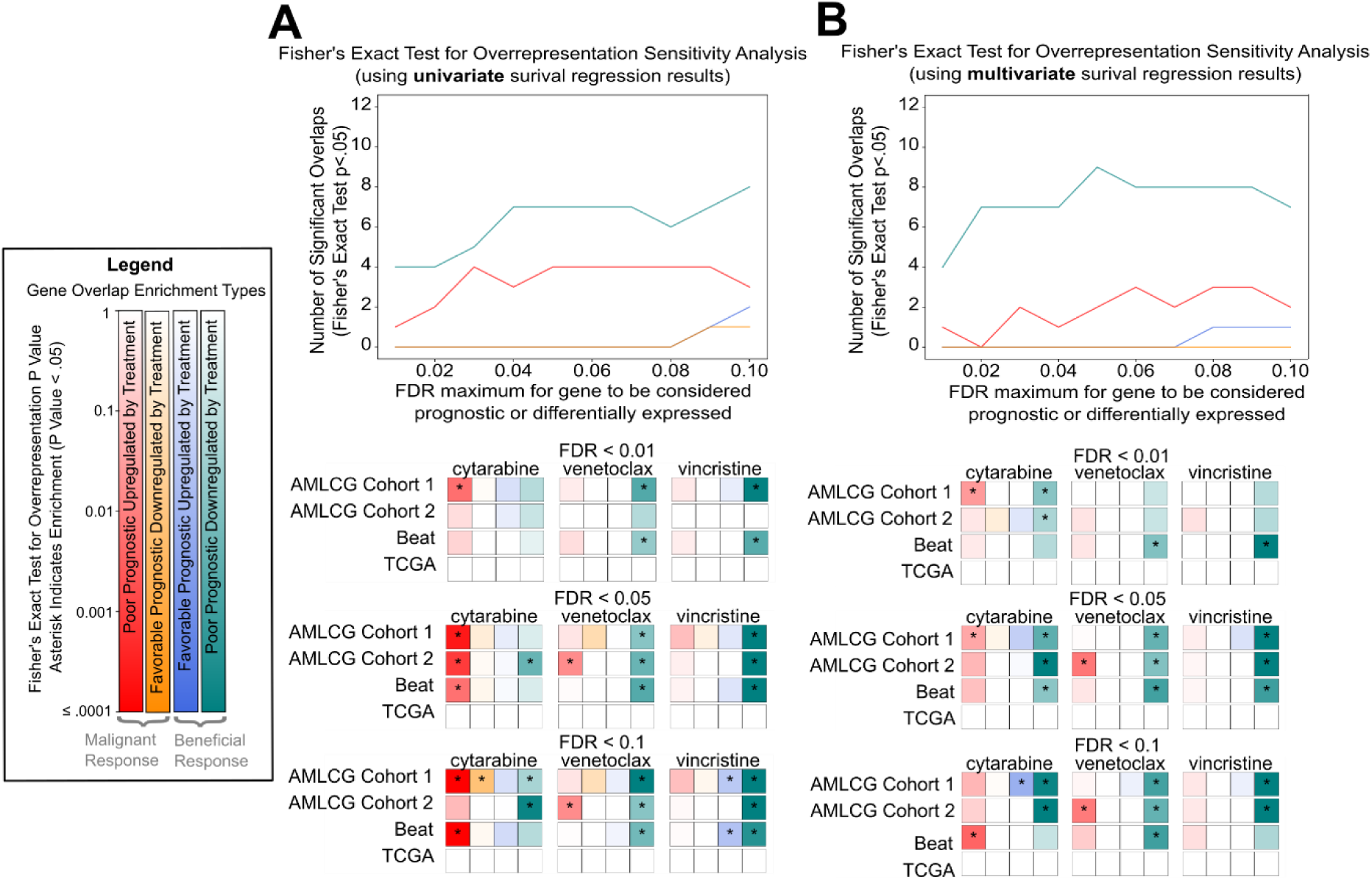
Sensitivity Analysis of Enrichment Testing Between Drug Induced and Prognostic Genes. Sensitivity analysis of enrichment results between drug induced and prognostic genes presented in Fig.1. Panels corresponds to the type of survival analysis (univariate or multivariate) used to determine gene prognosis associations. Within each panel, line plots at the top indicate the number of significant enrichment results (p < 0.05, y-axis) for each type of enrichment (line color) and how this varies based on a shared FDR stringency (x-axis) for classifying genes as prognostic or differentially expressed (provided log2FC > .7), assessed between 0.01 and 0.1 at increments of 0.01. Below, grid plots provide more detailed results corresponding to the FDR stringencies of .01, .05, and .1 along the x-axes of above line plots. For each stringency, 12 horizontal bars (one for each combination of drug and cohort choice) are shown, each bar containing 4 distinctly colored squares for each of the different enrichment types (poor prognostic and upregulated, favorable prognostic and down regulated, favorable prognostic and upregulated, poor prognostic and downregulated). Statistical enrichment for an enrichment type (Fisher’s exact test p value < .05) is denoted by an asterisk **A)** Sensitivity analysis using Fisher’s exact test for overrepresentation with univariate survival analysis results. **B)** Sensitivity analysis using Fisher’s exact test for overrepresentation with multivariate survival analysis results.

**Supplementary Figure 7:**
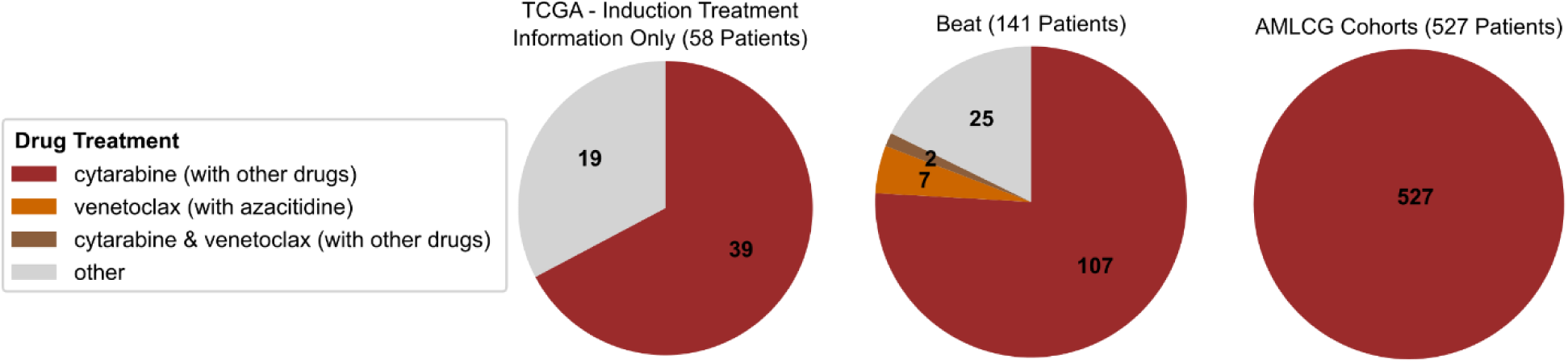
Drug Treatments Received by Survival Analysis Cohort Patients. Drug treatments administered to patients, with wedges indicating the proportion (and number) of patients receiving a given treatment in each cohort. Wedge colors distinguish drugs from our HL-60 experiments that were administered in cohorts (cytarabine, venetoclax, both, or neither), noting that these drugs were always given combinatorially, sometimes with drugs not administered in HL-60. TCGA data is only provided for induction treatment and does not explicitly label drugs, but cytarabine is understood in any “7+3” label to indicate 7 days of cytarabine with 3 days of an anthracycline as is common in abbreviating standard of care treatments. All documented treatment information can be found as specified in methods section under “data availability”.

## References

1. Sharma SV, Lee DY, Li B, Quinlan MP, Takahashi F, Maheswaran S, et al. A chromatin-mediated reversible drug-tolerant state in cancer cell subpopulations. Cell. 2010 Apr 2;141(1):69–80.

2. Pisco AO, Brock A, Zhou J, Moor A, Mojtahedi M, Jackson D, et al. Non-Darwinian dynamics in therapy-induced cancer drug resistance. Nat Commun. 2013;4:2467.

3. Hoek KS, Goding CR. Cancer stem cells versus phenotype-switching in melanoma. Pigment Cell Melanoma Res. 2010 Dec;23(6):746–59.

4. Vlashi E, Pajonk F. Cancer stem cells, cancer cell plasticity and radiation therapy. Semin Cancer Biol. 2015 Apr;31:28–35.

5. Muthukrishnan SD, Kawaguchi R, Nair P, Prasad R, Qin Y, Johnson M, et al. P300 promotes tumor recurrence by regulating radiation-induced conversion of glioma stem cells to vascular-like cells. Nat Commun. 2022 Oct 19;13(1):6202.

6. Behranvand N, Nasri F, Zolfaghari Emameh R, Khani P, Hosseini A, Garssen J, et al. Chemotherapy: a double-edged sword in cancer treatment. Cancer Immunol Immunother CII. 2022 Mar;71(3):507–26.

7. Lu W, Kang Y. Epithelial-Mesenchymal Plasticity in Cancer Progression and Metastasis. Dev Cell. 2019 May 6;49(3):361–74.

8. Martins-Neves SR, Paiva-Oliveira DI, Wijers-Koster PM, Abrunhosa AJ, Fontes-Ribeiro C, Bovée JVMG, et al. Chemotherapy induces stemness in osteosarcoma cells through activation of Wnt/β - catenin signaling. Cancer Lett. 2016 Jan 28;370(2):286–95.

9. Dean M, Fojo T, Bates S. Tumour stem cells and drug resistance. Nat Rev Cancer. 2005 Apr;5(4):275–84.

10. Blanpain C, Mohrin M, Sotiropoulou PA, Passegué E. DNA-damage response in tissue-specific and cancer stem cells. Cell Stem Cell. 2011 Jan 7;8(1):16–29.

11. Sharma A, Jasrotia S, Kumar A. Effects of Chemotherapy on the Immune System: Implications for Cancer Treatment and Patient Outcomes. Naunyn Schmiedebergs Arch Pharmacol. 2024 May;397(5):2551–66.

12. Su Y, Wei W, Robert L, Xue M, Tsoi J, Garcia-Diaz A, et al. Single-cell analysis resolves the cell state transition and signaling dynamics associated with melanoma drug-induced resistance. Proc Natl Acad Sci U S A. 2017 Dec 26;114(52):13679–84.

13. Vyas D, Laput G, Vyas AK. Chemotherapy-enhanced inflammation may lead to the failure of therapy and metastasis. OncoTargets Ther. 2014;7:1015–23.

14. Deyell M, Garris CS, Laughney AM. Cancer metastasis as a non-healing wound. Br J Cancer. 2021 Apr;124(9):1491–502.

15. Testa U, Castelli G, Pelosi E. Recent Developments in Differentiation Therapy of Acute Myeloid Leukemia. Cancers. 2025 Mar 28;17(7):1141.

16. Huang AC, Hu L, Kauffman SA, Zhang W, Shmulevich I. Using cell fate attractors to uncover transcriptional regulation of HL60 neutrophil differentiation. BMC Syst Biol. 2009 Feb 18;3:20.

17. Zhu K, Xia Y, Tian X, He Y, Zhou J, Han R, et al. Characterization and therapeutic perspectives of differentiation-inducing therapy in malignant tumors. Front Genet. 2023 Sep 8;14:1271381.

18. van Gils N, Verhagen HJMP, Smit L. Reprogramming acute myeloid leukemia into sensitivity for retinoic-acid-driven differentiation. Exp Hematol. 2017 Aug 1;52:12–23.

19. Huang S. The Logic of Cancer Treatment: Why It Is So Hard to Cure Cancer; Treatment-Induced Progression, Hyper-Progression, and the Nietzsche Effect. In: Strauss B, Bertolaso M, Ernberg I, Bissell MJ, editors. Rethinking Cancer [Internet]. The MIT Press; 2021 [cited 2025 Aug 5]. p. 63–128. Available from: https://direct.mit.edu/books/book/5122/chapter/3099567/The-Logic-of-Cancer-Treatment-Why-It-Is-So-Hard-to

20. Huang S, Ernberg I, Kauffman S. Cancer attractors: a systems view of tumors from a gene network dynamics and developmental perspective. Semin Cell Dev Biol. 2009 Sep;20(7):869–76.

21. Shin D, Gong JR, Jeong SD, Cho Y, Kim HP, Kim TY, et al. Attractor Landscape Analysis Reveals a Reversion Switch in the Transition of Colorectal Tumorigenesis. Adv Sci Weinh Baden-Wurtt Ger. 2025 Feb;12(8):e2412503.

22. Li Q, Wennborg A, Aurell E, Dekel E, Zou JZ, Xu Y, et al. Dynamics inside the cancer cell attractor reveal cell heterogeneity, limits of stability, and escape. Proc Natl Acad Sci U S A. 2016 Mar 8;113(10):2672–7.

23. Maetschke SR, Ragan MA. Characterizing cancer subtypes as attractors of Hopfield networks. Bioinforma Oxf Engl. 2014 May 1;30(9):1273–9.

24. Park JH, Hothi P, Lopez Garcia de Lomana A, Pan M, Calder R, Turkarslan S, et al. Gene regulatory network topology governs resistance and treatment escape in glioma stem-like cells. BioRxiv Prepr Serv Biol. 2024 Feb 7;2024.02.02.578510.

25. Bottomly D, Long N, Schultz AR, Kurtz SE, Tognon CE, Johnson K, et al. Integrative analysis of drug response and clinical outcome in acute myeloid leukemia. Cancer Cell. 2022 Aug;40(8):850–864.e9.

26. The Cancer Genome Atlas Research Network. Genomic and Epigenomic Landscapes of Adult De Novo Acute Myeloid Leukemia. N Engl J Med. 2013 May 30;368(22):2059–74.

27. Büchner T, Berdel WE, Schoch C, Haferlach T, Serve HL, Kienast J, et al. Double induction containing either two courses or one course of high-dose cytarabine plus mitoxantrone and postremission therapy by either autologous stem-cell transplantation or by prolonged maintenance for acute myeloid leukemia. J Clin Oncol Off J Am Soc Clin Oncol. 2006 Jun 1;24(16):2480–9.

28. Irizarry RA, Bolstad BM, Collin F, Cope LM, Hobbs B, Speed TP. Summaries of Affymetrix GeneChip probe level data. Nucleic Acids Res. 2003 Feb 15;31(4):e15.

29. Herold T, Jurinovic V, Metzeler KH, Boulesteix AL, Bergmann M, Seiler T, et al. An eight-gene expression signature for the prediction of survival and time to treatment in chronic lymphocytic leukemia. Leukemia. 2011 Oct;25(10):1639–45.

30. Li Z, Herold T, He C, Valk PJM, Chen P, Jurinovic V, et al. Identification of a 24-gene prognostic signature that improves the European LeukemiaNet risk classification of acute myeloid leukemia: an international collaborative study. J Clin Oncol Off J Am Soc Clin Oncol. 2013 Mar 20;31(9):1172–81.

31. Davidson-Pilon C. lifelines: survival analysis in Python. J Open Source Softw. 2019 Aug 4;4(40):1317.

32. Bray NL, Pimentel H, Melsted P, Pachter L. Near-optimal probabilistic RNA-seq quantification. Nat Biotechnol. 2016 May;34(5):525–7.

33. Love MI, Huber W, Anders S. Moderated estimation of fold change and dispersion for RNA-seq data with DESeq2. Genome Biol. 2014 Dec 5;15(12):550.

34. Benjamini Y, Hochberg Y. Controlling the False Discovery Rate: A Practical and Powerful Approach to Multiple Testing. J R Stat Soc Ser B Stat Methodol. 1995 Jan 1;57(1):289–300.

35. Melsted P, Booeshaghi AS, Liu L, Gao F, Lu L, Min KH, et al. Modular, efficient and constant-memory single-cell RNA-seq preprocessing. Nat Biotechnol. 2021 Jul;39(7):813–8.

36. Wolf FA, Angerer P, Theis FJ. SCANPY: large-scale single-cell gene expression data analysis. Genome Biol. 2018 Dec;19(1):15.

37. Chen EY, Tan CM, Kou Y, Duan Q, Wang Z, Meirelles GV, et al. Enrichr: interactive and collaborative HTML5 gene list enrichment analysis tool. BMC Bioinformatics. 2013 Apr 15;14:128.

38. Mehta CR, Patel NR. A Network Algorithm for Performing Fisher’s Exact Test in r × c Contingency Tables. J Am Stat Assoc. 1983 Jun;78(382):427.

39. Mehta CR, Patel NR. ALGORITHM 643: FEXACT: a FORTRAN subroutine for Fisher’s exact test on unordered *r×c* contingency tables. ACM Trans Math Softw. 1986 Jun;12(2):154–61.

40. Keenan AB, Torre D, Lachmann A, Leong AK, Wojciechowicz ML, Utti V, et al. ChEA3: transcription factor enrichment analysis by orthogonal omics integration. Nucleic Acids Res. 2019 Jul 2;47(W1):W212–24.

41. Ohno R, Kobayashi T, Tanimoto M, Hiraoka A, Imai K, Asou N, et al. Randomized study of individualized induction therapy with or without vincristine, and of maintenance-intensification therapy between 4 or 12 courses in adult acute myeloid leukemia. AML-87 Study of the Japan Adult Leukemia Study Group. Cancer. 1993 Jun 15;71(12):3888–95.

42. Seegars MB, Woods R, Ellis LR, Bhave RR, Howard DS, Manuel M, et al. A Pilot Phase II Study of the Feasibility and Efficacy of Vincristine Sulfate Liposome Injection in Patients With Relapsed or Refractory Acute Myeloid Leukemia. J Hematol. 2021 Feb;10(1):1–7.

43. Whiting FJH, Mossner M, Gabbutt C, Kimberley C, Barnes CP, Baker AM, et al. Quantitative measurement of phenotype dynamics during cancer drug resistance evolution using genetic barcoding. Nat Commun. 2025 Jun 20;16(1):5282.

44. Shen S, Clairambault J. Cell plasticity in cancer cell populations. F1000Research. 2020;9:F1000 Faculty Rev–635.

45. Chang HH, Hemberg M, Barahona M, Ingber DE, Huang S. Transcriptome-wide noise controls lineage choice in mammalian progenitor cells. Nature. 2008 May 22;453(7194):544–7.

46. Gardner AL, Jost TA, Morgan D, Brock A. Computational identification of surface markers for isolating distinct subpopulations from heterogeneous cancer cell populations. NPJ Syst Biol Appl. 2024 Oct 17;10(1):120.

47. Demaria M, O’Leary MN, Chang J, Shao L, Liu S, Alimirah F, et al. Cellular Senescence Promotes Adverse Effects of Chemotherapy and Cancer Relapse. Cancer Discov. 2017 Feb;7(2):165–76.

48. Angelini E, Wang Y, Zhou JX, Qian H, Huang S. A model for the intrinsic limit of cancer therapy: Duality of treatment-induced cell death and treatment-induced stemness. Alber M, editor. PLOS Comput Biol. 2022 Jul 25;18(7):e1010319.

49. Howard GR, Jost TA, Yankeelov TE, Brock A. Quantification of long-term doxorubicin response dynamics in breast cancer cell lines to direct treatment schedules. PLoS Comput Biol. 2022 Mar;18(3):e1009104.

50. Theile D, Wizgall P. Acquired ABC-transporter overexpression in cancer cells: transcriptional induction or Darwinian selection? Naunyn Schmiedebergs Arch Pharmacol. 2021 Aug;394(8):1621–32.

51. Johnson KE, Howard GR, Morgan D, Brenner EA, Gardner AL, Durrett RE, et al. Integrating transcriptomics and bulk time course data into a mathematical framework to describe and predict therapeutic resistance in cancer. Phys Biol. 2020 Nov 20;18(1):016001.

52. Qin S, Jiang J, Lu Y, Nice EC, Huang C, Zhang J, et al. Emerging role of tumor cell plasticity in modifying therapeutic response. Signal Transduct Target Ther. 2020 Oct 7;5(1):228.

53. Haak VM, Huang S, Panigrahy D. Debris-stimulated tumor growth: a Pandora’s box? Cancer Metastasis Rev. 2021 Sep;40(3):791–801.

54. Sulciner ML, Serhan CN, Gilligan MM, Mudge DK, Chang J, Gartung A, et al. Resolvins suppress tumor growth and enhance cancer therapy. J Exp Med. 2018 Jan 2;215(1):115–40.

55. Revesz L. Effect of tumour cells killed by x-rays upon the growth of admixed viable cells. Nature. 1956 Dec 22;178(4547):1391–2.

56. Willems JJLP, Arnold BP, Gregory CD. Sinister self-sacrifice: the contribution of apoptosis to malignancy. Front Immunol. 2014;5:299.

57. Morana O, Wood W, Gregory CD. The Apoptosis Paradox in Cancer. Int J Mol Sci. 2022 Jan 25;23(3):1328.

58. Kantarjian H, Kadia T, DiNardo C, Daver N, Borthakur G, Jabbour E, et al. Acute myeloid leukemia: current progress and future directions. Blood Cancer J [Internet]. 2021 Feb 22 [cited 2025 May 23];11(2). Available from: https://www.nature.com/articles/s41408-021-00425-3

59. Döhner H, Wei AH, Appelbaum FR, Craddock C, DiNardo CD, Dombret H, et al. Diagnosis and management of AML in adults: 2022 recommendations from an international expert panel on behalf of the ELN. Blood. 2022 Sep 22;140(12):1345–77.

60. Tang H, Wang S, Xiao G, Schiller J, Papadimitrakopoulou V, Minna J, et al. Comprehensive evaluation of published gene expression prognostic signatures for biomarker-based lung cancer clinical studies. Ann Oncol Off J Eur Soc Med Oncol. 2017 Apr 1;28(4):733–40.

61. Hukku A, Quick C, Luca F, Pique-Regi R, Wen X. BAGSE: a Bayesian hierarchical model approach for gene set enrichment analysis. Hancock J, editor. Bioinformatics. 2020 Mar 1;36(6):1689–95.

62. Smith KR, Slattery ML, French TK. Collinear nutrients and the risk of colon cancer. J Clin Epidemiol. 1991;44(7):715–23.

63. Domburg RT van, Hoeks SE, Kardys I, Lenzen M, Boersma E. Tools and Techniques - Statistics: How many variables are allowed in the logistic and Cox regression models? [Internet]. [cited 2024 Dec 19]. Available from: https://eurointervention.pcronline.com/article/tools-and-techniques-statistics-how-many-variables-are-allowed-in-the-logistic-and-cox-regression-models

64. Xue X, Kim MY, Shore RE. Cox regression analysis in presence of collinearity: an application to assessment of health risks associated with occupational radiation exposure. Lifetime Data Anal. 2007 Sep 1;13(3):333–50.

65. Concato J, Feinstein AR, Holford TR. The Risk of Determining Risk with Multivariable Models. Ann Intern Med. 1993 Feb 1;118(3):201–10.

66. Singh S, Sarkar T, Jakubison B, Gadomski S, Spradlin A, Gudmundsson KO, et al. Inhibitor of DNA binding proteins revealed as orchestrators of steady state, stress and malignant hematopoiesis. Front Immunol. 2022 Aug 5;13:934624.

67. Fei MY, Wang Y, Chang BH, Xue K, Dong F, Huang D, et al. The non-cell-autonomous function of ID1 promotes AML progression via ANGPTL7 from the microenvironment. Blood. 2023 Sep 7;142(10):903–17.

68. Id1 restrains myeloid commitment, maintaining the self-renewal capacity of hematopoietic stem cells [Internet]. [cited 2024 Dec 19]. Available from: https://www.pnas.org/doi/epdf/10.1073/pnas.0607894104

69. Wang L, Man N, Sun XJ, Tan Y, Cao MG, Liu F, et al. Regulation of AKT signaling by Id1 controls t(8;21) leukemia initiation and progression. Blood. 2015 Jul 30;126(5):640–50.

70. Suh HC, Leeanansaksiri W, Ji M, Klarmann KD, Renn K, Gooya J, et al. Id1 immortalizes hematopoietic progenitors in vitro and promotes a myeloproliferative disease in vivo. Oncogene. 2008 Sep;27(42):5612–23.

71. Nigten J, Breems-de Ridder MC, Erpelinck-Verschueren CAJ, Nikoloski G, Van Der Reijden BA, Van Wageningen S, et al. ID1 and ID2 are retinoic acid responsive genes and induce a G0/G1 accumulation in acute promyelocytic leukemia cells. Leukemia. 2005 May 1;19(5):799–805.

72. Lima AS, Bezerra MF, Moreira-Aguiar A, Weinhäuser I, Santos BL, Falcão RM, et al. Prognostic implications of the ID1 expression in acute myeloid leukemia patients treated in a resource - constrained setting. Hematol Transfus Cell Ther. 2024;46(3):250–5.

73. Tang R, Hirsch P, Fava F, Lapusan S, Marzac C, Teyssandier I, et al. High Id1 expression is associated with poor prognosis in 237 patients with acute myeloid leukemia. Blood. 2009 Oct 1;114(14):2993–3000.

74. Zhou JD, Ma JC, Zhang TJ, Li XX, Zhang W, Wu DH, et al. High bone marrow ID2 expression predicts poor chemotherapy response and prognosis in acute myeloid leukemia. Oncotarget. 2017 Sep 1;8(54):91979–89.

75. Russell RG, Lasorella A, Dettin LE, Iavarone A. Id2 drives differentiation and suppresses tumor formation in the intestinal epithelium. Cancer Res. 2004 Oct 15;64(20):7220–5.

76. Ghisi M, Kats L, Masson F, Li J, Kratina T, Vidacs E, et al. Id2 and E Proteins Orchestrate the Initiation and Maintenance of MLL-Rearranged Acute Myeloid Leukemia. Cancer Cell. 2016 Jul 11;30(1):59–74.

77. Wågsäter D, Sirsjö A, Dimberg J. Down-regulation of ID2 by all-trans retinoic acid in monocytic leukemia cells (THP-1). J Exp Clin Cancer Res CR. 2003 Sep;22(3):471–5.

78. May AM, Frey AV, Bogatyreva L, Benkisser-Petersen M, Hauschke D, Lübbert M, et al. ID2 and ID3 protein expression mirrors granulopoietic maturation and discriminates between acute leukemia subtypes. Histochem Cell Biol. 2014 Apr;141(4):431–40.

79. Benetatos L, Dasoula A, Hatzimichael E, Syed N, Voukelatou M, Dranitsaris G, et al. Polo-like kinase 2 (SNK/PLK2) is a novel epigenetically regulated gene in acute myeloid leukemia and myelodysplastic syndromes: genetic and epigenetic interactions. Ann Hematol. 2011 Sep 1;90(9):1037–45.

80. Burns TF, Fei P, Scata KA, Dicker DT, El-Deiry WS. Silencing of the novel p53 target gene Snk/Plk2 leads to mitotic catastrophe in paclitaxel (taxol)-exposed cells. Mol Cell Biol. 2003 Aug;23(16):5556–71.

81. Schiera G, Di Liegro CM, Puleo V, Colletta O, Fricano A, Cancemi P, et al. Extracellular vesicles shed by melanoma cells contain a modified form of H1.0 linker histone and H1.0 mRNA-binding proteins. Int J Oncol. 2016 Nov;49(5):1807–14.

82. Schiera G, Di Liegro CM, Saladino P, Pitti R, Savettieri G, Proia P, et al. Oligodendroglioma cells synthesize the differentiation-specific linker histone H1° and release it into the extracellular environment through shed vesicles. Int J Oncol. 2013 Dec;43(6):1771–6.

83. Lord KA, Abdollahi A, Hoffman-Liebermann B, Liebermann DA. Dissection of the immediate early response of myeloid leukemia cells to terminal differentiation and growth inhibitory stimuli. Cell Growth Differ Mol Biol J Am Assoc Cancer Res. 1990 Dec;1(12):637–45.

84. Kohli A, Huang SL, Chang TC, Chao CCK, Sun NK. H1.0 induces paclitaxel-resistance genes expression in ovarian cancer cells by recruiting GCN5 and androgen receptor. Cancer Sci. 2022 Aug;113(8):2616–26.

85. Morales Torres C, Wu MY, Hobor S, Wainwright EN, Martin MJ, Patel H, et al. Selective inhibition of cancer cell self-renewal through a Quisinostat-histone H1.0 axis. Nat Commun. 2020 Apr 14;11(1):1792.

86. Torres CM, Biran A, Burney MJ, Patel H, Henser-Brownhill T, Cohen AHS, et al. The linker histone H1.0 generates epigenetic and functional intratumor heterogeneity. Science. 2016 Sep 30;353(6307):aaf1644.

87. Mii S, Enomoto A, Shiraki Y, Taki T, Murakumo Y, Takahashi M. CD109: a multifunctional GPI-anchored protein with key roles in tumor progression and physiological homeostasis. Pathol Int. 2019 May;69(5):249–59.

88. CD109 is expressed on a subpopulation of CD34+ cells enriched in hematopoietic stem and progenitor cells - PubMed [Internet]. [cited 2025 Jan 16]. Available from: https://pubmed.ncbi.nlm.nih.gov/10428505/

89. The novel prognostic analysis of AML based on ferroptosis and cuproptosis related genes - ScienceDirect [Internet]. [cited 2025 Jan 16]. Available from: https://www.sciencedirect.com/science/article/pii/S0946672X24001378?via%3Dihub#sec0010

90. Mining of transcriptome identifies CD109 and LRP12 as possible biomarkers and deregulation mechanism of T cell receptor pathway in Acute Myeloid Leukemia - PMC [Internet]. [cited 2025 Jan 16]. Available from: https://pmc.ncbi.nlm.nih.gov/articles/PMC9589179/#sec5

91. A parsimonious 3-gene signature predicts clinical outcomes in an acute myeloid leukemia multicohort study - PubMed [Internet]. [cited 2025 Jan 16]. Available from: https://pubmed.ncbi.nlm.nih.gov/31015209/

92. Duan C, Lin X, Zou W, He Q, Wei F, Pan J, et al. Targeting DDX3X eliminates leukemia stem cells in chronic myeloid leukemia by blocking NT5DC2 mRNA translation. Oncogene. 2025 Feb;44(4):241–54.

93. Li KS, Zhu XD, Liu HD, Zhang SZ, Li XL, Xiao N, et al. NT5DC2 promotes tumor cell proliferation by stabilizing EGFR in hepatocellular carcinoma. Cell Death Dis. 2020 May 7;11(5):335.

94. Sha R, Zhang J, Meng F, Zhaori G. Gastric cancer metastasis-related NT5DC2 indicates unfavorable prognosis of patients. Medicine (Baltimore). 2023 Oct 6;102(40):e35030.

95. Zhu Z, Hou Q, Guo H. NT5DC2 knockdown inhibits colorectal carcinoma progression by repressing metastasis, angiogenesis and tumor-associated macrophage recruitment: A mechanism involving VEGF signaling. Exp Cell Res. 2020 Dec 1;397(1):112311.

96. Cuevas D, Amigo R, Agurto A, Heredia AA, Guzmán C, Recabal-Beyer A, et al. The Role of Epithelial-to-Mesenchymal Transition Transcription Factors (EMT-TFs) in Acute Myeloid Leukemia Progression. Biomedicines. 2024 Aug 21;12(8):1915.

97. Gouda MBY, Hassan NM, Kandil EI. Bone marrow overexpression of SNAI1 is an early indicator of intrinsic drug resistance in patients with de novo acute myeloid leukemia. J Gene Med. 2023 May;25(5):e3443.

98. The EMT modulator SNAI1 contributes to AML pathogenesis via its interaction with LSD1 - PubMed [Internet]. [cited 2025 Jan 16]. Available from: https://pubmed.ncbi.nlm.nih.gov/32369597/

99. Saarikoski ST, Rivera SP, Hankinson O, Husgafvel-Pursiainen K. CYP2S1: A short review. Toxicol Appl Pharmacol. 2005 Sep 1;207(2, Supplement):62–9.

100. Smith G, Wolf CR, Deeni YY, Dawe RS, Evans AT, Comrie MM, et al. Cutaneous expression of cytochrome P450 CYP2S1: individuality in regulation by therapeutic agents for psoriasis and other skin diseases. The Lancet. 2003 Apr 19;361(9366):1336–43.

101. Cozzolino F, Rubartelli A, Aldinucci D, Sitia R, Torcia M, Shaw A, et al. Interleukin 1 as an autocrine growth factor for acute myeloid leukemia cells. Proc Natl Acad Sci U S A. 1989 Apr;86(7):2369–73.

102. Carey A, Edwards DK, Eide CA, Newell L, Traer E, Medeiros BC, et al. Identification of interleukin-1 by functional screening as a key mediator of cellular expansion and disease progression in acute myeloid leukemia. Cell Rep. 2017 Mar 28;18(13):3204–18.

103. Sakai K, Hattori T, Matsuoka M, Asou N, Yamamoto S, Sagawa K, et al. Autocrine stimulation of interleukin 1 beta in acute myelogenous leukemia cells. J Exp Med. 1987 Nov 1;166(5):1597–602.

104. Rabe JL, Gardner L, Hunter R, Fonseca JA, Dougan J, Gearheart CM, et al. IL12 Abrogates Calcineurin-Dependent Immune Evasion during Leukemia Progression. Cancer Res. 2019 Jul 15;79(14):3702–13.

105. Landoni E, Woodcock MG, Barragan G, Casirati G, Cinella V, Stucchi S, et al. IL-12 reprograms CAR-expressing natural killer T cells to long-lived Th1-polarized cells with potent antitumor activity. Nat Commun. 2024 Jan 2;15(1):89.

106. Ferretti E, Cocco C, Airoldi I, Pistoia V. Targeting acute myeloid leukemia cells with cytokines. J Leukoc Biol. 2012 Sep;92(3):567–75.

107. Dunussi-Joannopoulos K, Leonard JP. Interleukin-12 gene therapy vaccines: directing the immune system against minimal residual leukemia. Leuk Lymphoma. 2001 May;41(5–6):483–92.

108. Bai J, Zhao Y, Shi K, Fan Y, Ha Y, Chen Y, et al. HIF-1α-mediated LAMC1 overexpression is an unfavorable predictor of prognosis for glioma patients: evidence from pan-cancer analysis and validation experiments. J Transl Med. 2024 Apr 27;22(1):391.

109. Liu J, Liu D, Yang Z, Yang Z. High LAMC1 expression in glioma is associated with poor prognosis. OncoTargets Ther. 2019;12:4253–60.

110. Ye G, Qin Y, Wang S, Pan D, Xu S, Wu C, et al. Lamc1 promotes the Warburg effect in hepatocellular carcinoma cells by regulating PKM2 expression through AKT pathway. Cancer Biol Ther. 2019;20(5):711–9.

111. Nishikawa R, Goto Y, Kojima S, Enokida H, Chiyomaru T, Kinoshita T, et al. Tumor-suppressive microRNA-29s inhibit cancer cell migration and invasion via targeting LAMC1 in prostate cancer. Int J Oncol. 2014 Jul;45(1):401–10.

112. Karabay AZ, Ozkan T, Karadag Gurel A, Koc A, Hekmatshoar Y, Sunguroglu A, et al. Identification of exosomal microRNAs and related hub genes associated with imatinib resistance in chronic myeloid leukemia. Naunyn Schmiedebergs Arch Pharmacol. 2024;397(12):9701–21.

113. Zhang J, Zhang M, Liang Y, Liu M, Huang Z. Downregulation of Smad4 expression confers chemoresistance against imatinib mesylate to chronic myeloid leukemia K562 cells. Hematol Amst Neth. 2022 Dec;27(1):43–52.

114. Yang LH, Wang GZ, Gao C. SEPT3 as a Potential Molecular Target of Triple-Negative Breast Cancer. Int J Gen Med. 2024;17:1605–13.

115. Ragaini S, Wagner S, Marconi G, Parisi S, Sartor C, Nanni J, et al. An IDO1-related immune gene signature predicts overall survival in acute myeloid leukemia. Blood Adv. 2022 Jan 11;6(1):87–99.

116. Qin L, Li B, Wang S, Tang Y, Fahira A, Kou Y, et al. Construction of an immune - related prognostic signature and lncRNA-miRNA-mRNA ceRNA network in acute myeloid leukemia. J Leukoc Biol. 2024 Jun 28;116(1):146–65.

117. König K, Marth L, Roissant J, Granja T, Jennewein C, Devanathan V, et al. The plexin C1 receptor promotes acute inflammation. Eur J Immunol. 2014 Sep;44(9):2648–58.

118. Abegunde SO, Grieve S, Reiman T. TAZ downregulated ANXA1 expression to modulate myeloma cell interactions with bone marrow mesenchymal stromal cells. Exp Hematol. 2024 Oct;138:104282.

119. Kotepui KU, Kotepui M, Piwkham D, Songsri A, Charoenkijkajorn L, Kongnok T, et al. Tissue Expression Of LPHN3 in Breast Cancer: An Immunohistochemistry Method. Asian Pac J Cancer Prev APJCP. 2020 Nov 1;21(11):3339–43.

120. Goto T, Teramoto Y, Nagata Y, Miyamoto H. Latrophilin-3 as a downstream effector of the androgen receptor induces bladder cancer progression. Discov Oncol. 2024 Sep 13;15(1):440.

121. Li M, Hao S, Li C, Xiao H, Sun L, Yu Z, et al. Elevated SH3BP5 Correlates with Poor Outcome and Contributes to the Growth of Acute Myeloid Leukemia Cells. Biomolecules. 2019 Sep 19;9(9):505.

122. Wang X, Liu R, Wang Y, Cai H, Zhang L. Effects of down-regulation of clusterin by small interference RNA on human acute myeloid leukemia cells. Int J Clin Exp Med. 2015;8(11):20925–31.

123. Kim DY, Shin DY, Oh S, Kim I, Kim EJ. Gene Expression and DNA Methylation Profiling Suggest Potential Biomarkers for Azacitidine Resistance in Myelodysplastic Syndrome. Int J Mol Sci. 2024 Apr 26;25(9):4723.

124. Ozel B, Kipcak S, Biray Avci C, Sabour Takanlou M, Sabour Takanlou L, Tezcanli Kaymaz B, et al. Targeting UPR signaling pathway by dasatinib as a promising therapeutic approach in chronic myeloid leukemia. Med Oncol Northwood Lond Engl. 2022 Jun 18;39(9):126.

125. Xia L min, Tian D an, Zhang Q, Yan W, Wang B, Liu M, et al. [Inhibition of HSP70-2 expression by RNA interference induces apoptosis of human hepatocellular carcinoma cells]. Zhonghua Gan Zang Bing Za Zhi Zhonghua Ganzangbing Zazhi Chin J Hepatol. 2008 Sep;16(9):678–82.

126. Radhakrishnan K, Truong L, Carmichael CL. An “unexpected” role for EMT transcription factors in hematological development and malignancy. Front Immunol [Internet]. 2023 Aug 3 [cited 2024 Dec 20];14. Available from: https://www.frontiersin.org/journals/immunology/articles/10.3389/fimmu.2023.1207360/full

127. Quintero-Ronderos P, Laissue P. The multisystemic functions of FOXD1 in development and disease. J Mol Med Berl Ger. 2018 Aug;96(8):725–39.

128. Zhao YF, Zhao JY, Yue H, Hu KS, Shen H, Guo ZG, et al. FOXD1 promotes breast cancer proliferation and chemotherapeutic drug resistance by targeting p27. Biochem Biophys Res Commun. 2015 Jan 2;456(1):232–7.

129. Cai K, Chen S, Zhu C, Li L, Yu C, He Z, et al. FOXD1 facilitates pancreatic cancer cell proliferation, invasion, and metastasis by regulating GLUT1-mediated aerobic glycolysis. Cell Death Dis. 2022 Sep 3;13(9):765.

130. Qiu J, Li M, Su C, Liang Y, Ou R, Chen X, et al. FOXS1 Promotes Tumor Progression by Upregulating CXCL8 in Colorectal Cancer. Front Oncol. 2022;12:894043.

131. Wang M, Huang W. FOXS1 promotes prostate cancer progression through the Hedgehog/Gli1 pathway. Biochem Pharmacol. 2023 Dec;218:115893.

132. Ren R, Wang H, Xu Y, Wu J, Ma D, Guan W. FOXS1 acts as an oncogene and induces EMT through FAK/PI3K/AKT pathway by upregulating HILPDA in prostate cancer. FASEB J Off Publ Fed Am Soc Exp Biol. 2024 May 31;38(10):e23698.

133. Machon O, Masek J, Machonova O, Krauss S, Kozmik Z. Meis2 is essential for cranial and cardiac neural crest development. BMC Dev Biol. 2015 Nov 6;15:40.

134. Vegi NM, Klappacher J, Oswald F, Mulaw MA, Mandoli A, Thiel VN, et al. MEIS2 Is an Oncogenic Partner in AML1-ETO-Positive AML. Cell Rep. 2016 Jul;16(2):498–507.

135. Xiao Y, Liu Y, Sun Y, Huang C, Zhong S. MEIS2 suppresses breast cancer development by downregulating IL10. Cancer Rep Hoboken NJ. 2024 May;7(5):e2064.

136. Wang M, Wang H, Wen Y, Chen X, Liu X, Gao J, et al. MEIS2 regulates endothelial to hematopoietic transition of human embryonic stem cells by targeting TAL1. Stem Cell Res Ther. 2018 Dec 7;9(1):340.

137. Wen X, Liu M, Du J, Wang X. Meis homeobox 2 (MEIS2) inhibits the proliferation and promotes apoptosis of thyroid cancer cell and through the NF-κB signaling pathway. Bioengineered. 2021 Dec;12(1):1766–72.

138. Wan Z, Chai R, Yuan H, Chen B, Dong Q, Zheng B, et al. MEIS2 promotes cell migration and invasion in colorectal cancer. Oncol Rep. 2019 Jul;42(1):213–23.

139. Nørgaard M, Haldrup C, Bjerre MT, Høyer S, Ulhøi B, Borre M, et al. Epigenetic silencing of MEIS2 in prostate cancer recurrence. Clin Epigenetics. 2019 Oct 22;11(1):147.

140. You Y, Ma Y, Wang Q, Ye Z, Deng Y, Bai F. Attenuated ZHX3 expression serves as a potential biomarker that predicts poor clinical outcomes in breast cancer patients. Cancer Manag Res. 2019;11:1199–210.

141. Cai Z, Wang S, Zhou H, Cao D. Low expression of ZHX3 is associated with progression and poor prognosis in colorectal cancer. Transl Oncol. 2024 Jan;39:101829.

142. Deng M, Wei W, Duan J, Chen R, Wang N, He L, et al. ZHX3 promotes the progression of urothelial carcinoma of the bladder via repressing of RGS2 and is a novel substrate of TRIM21. Cancer Sci. 2021 May;112(5):1758–71.

143. Igata T, Tanaka H, Etoh K, Hong S, Tani N, Koga T, et al. Loss of the transcription repressor ZHX3 induces senescence-associated gene expression and mitochondrial-nucleolar activation. PloS One. 2022;17(1):e0262488.

144. Feng Y, Zhang T, Wang Y, Xie M, Ji X, Luo X, et al. Homeobox Genes in Cancers: From Carcinogenesis to Recent Therapeutic Intervention. Front Oncol. 2021 Oct 14;11:770428.

145. Yang Y, Zhang M, Zhao Y, Deng T, Zhou X, Qian H, et al. HOXD8 suppresses renal cell carcinoma growth by upregulating SHMT1 expression. Cancer Sci. 2023 Dec;114(12):4583–95.

146. Yu K, Meng J, Chen T, Wang Y, Zhao Y, Huang T, et al. HOXD8 drives Glioma progression through epithelial-mesenchymal transition regulation: Implications for prognosis and targeted therapy. Exp Cell Res. 2025 Mar 15;446(2):114476.

147. Zhang Y, Yu Y, Su X, Lu Y. HOXD8 inhibits the proliferation and migration of triple - negative breast cancer cells and induces apoptosis in them through regulation of AKT/mTOR pathway. Reprod Biol. 2021 Dec;21(4):100544.

148. Ran Y, Hu C, Wan J, Kang Q, Zhou R, Liu P, et al. Integrated investigation and experimental validation of PPARG as an oncogenic driver: implications for prognostic assessment and therapeutic targeting in hepatocellular carcinoma. Front Pharmacol. 2023;14:1298341.

149. Li DH, Liu XK, Tian XT, Liu F, Yao XJ, Dong JF. PPARG: A Promising Therapeutic Target in Breast Cancer and Regulation by Natural Drugs. PPAR Res. 2023;2023:4481354.

150. Tate T, Xiang T, Wobker SE, Zhou M, Chen X, Kim H, et al. Pparg signaling controls bladder cancer subtype and immune exclusion. Nat Commun. 2021 Oct 25;12(1):6160.

151. Villa ALP, Parra RS, Feitosa MR, Camargo HP de, Machado VF, Tirapelli DP da C, et al. PPARG expression in colorectal cancer and its association with staging and clinical evolution. Acta Cir Bras. 2020;35(7):e202000708.

152. Choi JH, Park JD, Choi SH, Ko ES, Jang HJ, Park KS. ELK3-ID4 axis governs the metastatic features of triple negative breast cancer. Oncol Res. 2023;32(1):127–38.

153. Lee M, Cho HJ, Park KS, Jung HY. ELK3 Controls Gastric Cancer Cell Migration and Invasion by Regulating ECM Remodeling-Related Genes. Int J Mol Sci. 2022 Mar 28;23(7):3709.

154. Mei Y, Chen D, He S, Ye J, Luo M, Wu Q, et al. Transcription Factor ELK3 Promotes Stemness and Oxaliplatin Resistance of Glioma Cells by Regulating RNASEH2A. Horm Metab Res Horm Stoffwechselforschung Horm Metab. 2023 Feb;55(2):149–55.

155. Meng L, Xing Z, Guo Z, Liu Z. LINC01106 post-transcriptionally regulates ELK3 and HOXD8 to promote bladder cancer progression. Cell Death Dis. 2020 Dec 12;11(12):1063.

156. Zhao Q, Ren Y, Xie H, Yu L, Lu J, Jiang W, et al. ELK3 Mediated by ZEB1 Facilitates the Growth and Metastasis of Pancreatic Carcinoma by Activating the Wnt/β-Catenin Pathway. Front Cell Dev Biol. 2021;9:700192.

157. Liao Y, Luo Z, Lin Y, Chen H, Chen T, Xu L, et al. PRMT3 drives glioblastoma progression by enhancing HIF1A and glycolytic metabolism. Cell Death Dis. 2022 Nov 9;13(11):943.

158. Qi H, Ma X, Ma Y, Jia L, Liu K, Wang H. Mechanisms of HIF1A-mediated immune evasion in gastric cancer and the impact on therapy resistance. Cell Biol Toxicol. 2024 Oct 10;40(1):87.

159. Yang K, Zhang W, Zhong L, Xiao Y, Sahoo S, Fassan M, et al. Long non-coding RNA HIF1A-As2 and MYC form a double-positive feedback loop to promote cell proliferation and metastasis in KRAS-driven non-small cell lung cancer. Cell Death Differ. 2023 Jun;30(6):1533–49.

160. Shih SP, Lu MC, El-Shazly M, Lin YH, Chen CL, Yu SSF, et al. The Antileukemic and Anti-Prostatic Effect of Aeroplysinin-1 Is Mediated through ROS-Induced Apoptosis via NOX Activation and Inhibition of HIF-1a Activity. Life Basel Switz. 2022 May 5;12(5):687.

161. Zhang Z, Shi J, Wu Q, Zhang Z, Liu X, Ren A, et al. JUN mediates glucocorticoid resistance by stabilizing HIF1a in T cell acute lymphoblastic leukemia. iScience. 2023 Nov 17;26(11):108242.

162. Kontos CK, Papageorgiou SG, Diamantopoulos MA, Scorilas A, Bazani E, Vasilatou D, et al. mRNA overexpression of the hypoxia inducible factor 1 alpha subunit gene (HIF1A): An independent predictor of poor overall survival in chronic lymphocytic leukemia. Leuk Res. 2017 Feb;53:65–73.

163. Chen J, Mu Q, Li X, Yin X, Yu M, Jin J, et al. Homoharringtonine targets Smad3 and TGF-β pathway to inhibit the proliferation of acute myeloid leukemia cells. Oncotarget. 2017 Jun 20;8(25):40318–26.

164. Runa F, Ortiz-Soto G, De Barros NR, Kelber JA. Targeting SMAD-Dependent Signaling: Considerations in Epithelial and Mesenchymal Solid Tumors. Pharmaceuticals. 2024 Mar 1;17(3):326.

165. Khan SF, Damerell V, Omar R, Du Toit M, Khan M, Maranyane HM, et al. The roles and regulation of TBX3 in development and disease. Gene. 2020 Feb 5;726:144223.

166. Lin KC, Park HW, Guan KL. Regulation of the Hippo Pathway Transcription Factor TEAD. Trends Biochem Sci. 2017 Nov;42(11):862–72.

167. An N, Peng H, Hou M, Su D, Wang L, Shen X, et al. The zinc figure protein ZNF575 impairs colorectal cancer growth via promoting p53 transcription. Oncol Res. 2023;31(3):307–16.

168. Xu F, Kong L, Sun X, Hui W, Jiang L, Han W, et al. PFDN6 contributes to colorectal cancer progression via transcriptional regulation. eGastroenterology. 2024 Apr;2(2):e100001.

169. Martínez-Aguilar L, Pérez-Ramírez C, Maldonado-Montoro MDM, Carrasco-Campos MI, Membrive-Jiménez C, Martínez-Martínez F, et al. Effect of genetic polymorphisms on therapeutic response in multiple sclerosis relapsing-remitting patients treated with interferon-beta. Mutat Res Rev Mutat Res. 2020;785:108322.

170. Correia JC, Jannig PR, Gosztyla ML, Cervenka I, Ducommun S, Præstholm SM, et al. Zfp697 is an RNA-binding protein that regulates skeletal muscle inflammation and remodeling. Proc Natl Acad Sci U S A. 2024 Aug 20;121(34):e2319724121.

171. Yu Z, Qiu B, Li L, Xu J, Zhou H, Niu T. An emerging prognosis prediction model for multiple myeloma: Hypoxia-immune related microenvironmental gene signature. Front Oncol. 2022;12:992387.

172. Ye M, Liu T, Miao L, Zou S, Ji H, Zhang J, et al. The Role of ZNF275/AKT Pathway in Carcinogenesis and Cisplatin Chemosensitivity of Cervical Cancer Using Patient-Derived Xenograft Models. Cancers. 2023 Nov 28;15(23):5625.

173. Yang W, Meyer AN, Jiang Z, Jiang X, Donoghue DJ. Critical domains for NACC2-NTRK2 fusion protein activation. Chen L, editor. PLOS ONE. 2024 Jun 27;19(6):e0301730.

174. Penning A, Snoeck S, Garritsen O, Tosoni G, Hof A, De Boer F, et al. NACC2, a molecular effector of miR-132 regulation at the interface between adult neurogenesis and Alzheimer’s disease. Sci Rep. 2024 Sep 10;14(1):21163.

